# Retrograde trafficking inhibitors allosterically trap Get3 to block tail-anchored protein biogenesis

**DOI:** 10.64898/2026.07.11.737968

**Authors:** Juliet A. Lee, Xilin Gu, Charlene Chan, Vida Storm Robertson, Victor Garcia-Ruiz, Yancheng E. Li, Anna Huyen Ngo, Philip Alabi, Vladimir Denic, Jason K. Sello, William M. Clemons

## Abstract

Retro-1 and Retro-2 are structurally distinct small molecules that protect cells from diverse toxins and viruses by disrupting retrograde trafficking, yet their mechanism of action has remained elusive. We show that both compounds target Get3, the ATPase chaperone of the guided entry of tail-anchored proteins (GET) pathway, which mediates biogenesis of tail-anchored SNARE proteins required for retrograde transport to the ER membrane. Cryo-electron microscopy reveals that Retro compounds bind a cryptic pocket in Get3, allosterically stabilizing Get3 in a stalled complex with upstream pathway components. Our work uncovers the GET pathway as an unsuspected vulnerability in pathogen entry, provides clear routes toward compound optimization, and establishes stabilization of dynamic protein complexes as a therapeutic strategy.

## Main Text

Within the secretory pathway, anterograde trafficking transports cargo from the endoplasmic reticulum (ER) through the Golgi to the cell surface, while retrograde trafficking returns cargo from endosomes through the Golgi to the ER. (*1*, *2*) (Fig. 1A). Diverse pathogens and toxins hijack host retrograde trafficking machinery to bypass lysosomal degradation and access their intracellular targets (*3*, *4*). Viruses, including coronaviruses, herpesviruses, and polyomaviruses, as well as the plant toxin ricin and bacterial Shiga-like toxins, depend on retrograde transport for cell entry and subsequent trafficking (*5–7*). Intracellular pathogens such as *Leishmania* and *Chlamydia* similarly exploit retrograde machinery to remodel host compartments to evade immune detection (*8*, *9*). Together, these convergent mechanisms of disease present retrograde trafficking as a therapeutic opportunity.

**Fig. 1.**
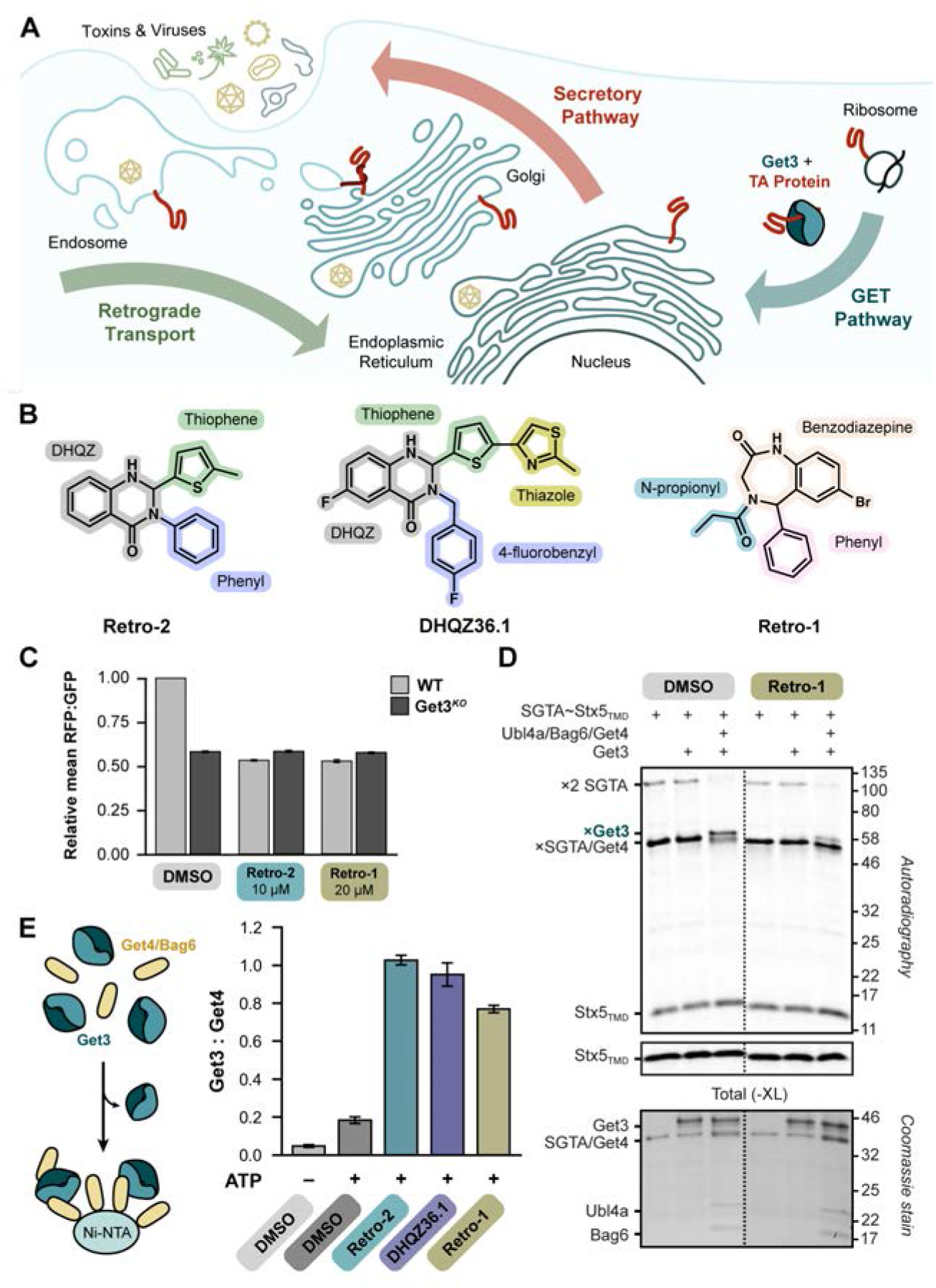
Retro compounds target Get3 and stabilize the pretargeting complex. (**A**) Schematic of the bidirectional secretory pathway highlighting retrograde toxin/virus transport (table S1). The GET pathway mediates biogenesis of TA proteins involved in both directions. Model TA protein shown in red. (**B**) Chemical structures of Retro-1, Retro-2, and DHQZ36.1 with key moieties labeled. Dihydroquinazolinone is abbreviated as DHQZ. (**C**) Wild-type (WT) and knockout (Get3^KO^) HEK293T cell lines with a GFP-2A-RFP-Sec61B_TMD_ reporter were treated as indicated and analyzed by fluorescence-activated cell sorting. RFP:GFP ratios were normalized to mock-treated WT (mean ± standard deviation, n = 3). DMSO- and Retro-2-treated data are from Morgens et al. 2019 (*27*). (**D**) ^35^S-methionine labeled SGTA-Stx5_TMD_ complex was incubated with indicated GET pathway components and Retro-1 or a DMSO control, crosslinked, and analyzed by SDS-PAGE with autoradiography (top) and Coomassie staining (bottom). Stx5_TMD_ adducts are marked with an (×). DMSO-treated data are from (*27*). (**E**) (left) Schematic of capture assay with Get3 dimer shown in teal and Get4/Bag6 complex shown in yellow. Get3–Get4/Bag6 capture assays were performed in the presence of indicated compounds. Coomassie-stained bands were quantified and shown as mean ± standard error.

In a search for compounds that protect against ricin toxicity, Retro-1 and Retro-2 were identified as selective, noncytotoxic inhibitors of retrograde transport from early endosomes to the trans-Golgi network (TGN) (*10*). Despite having distinct chemical scaffolds (*11–13*) (Fig. 1B), these compounds and their optimized analogs protect against a broad range of retrograde-trafficked toxins and pathogens (table S1) (*5–9*, *14–24*). Retro-2 has additionally been shown to disrupt autophagic vacuolar trafficking and inhibit breast cancer cell proliferation, underscoring the broader biological and medical relevance of these compounds (*25*, *26*).

Retro-2 treatment phenocopies the loss of components of the guided entry of tail-anchored proteins (GET) pathway, as revealed by a genome-wide screen (*27*). The GET pathway mediates post-translational targeting of tail-anchored (TA) proteins, membrane proteins with a single C-terminal transmembrane domain, to the ER (*28*, *29*), including SNARE proteins essential for vesicle fusion and retrograde transport (*30*, *31*). Notably, the broadly important target-SNARE Syntaxin-5 (Stx5) (*32*, *33*) is particularly sensitive to disruption of the GET pathway (*27*, *34*, *35*). While Retro-1 remains comparatively understudied, it also induces cell phenotypes resembling GET pathway mutations, including Stx5 mislocalization (*10*). These findings suggest that targeting the GET pathway could provide a broadly applicable strategy to treat infections, cancer, and toxin exposure (*26*, *35*, *36*).

Defining the precise mechanism by which Retro-2 targets the GET pathway has been historically challenging. Central to the GET pathway is the ATP-driven conformational cycling of the soluble ER-targeting factor Get3 between open and closed states (*37*). ATP-bound Get3 receives TA proteins from SGTA as part of a pretargeting complex that includes Get4, Ubl4a, and Bag6. TA protein handover triggers ATP hydrolysis and release of Get3 from the pretargeting complex (*38–40*). TA-bound Get3 is then recruited to the ER membrane by the WRB/CAML receptor for TA protein insertion (*41*, *42*). Retro-2 and its more potent analog, DHQZ36.1, were shown to block the transfer of newly synthesized TA proteins from SGTA to Get3. Critically, this effect is suppressed by point mutations in Get3 (*27*). Despite the body of work implicating Retro-2 as a GET pathway inhibitor, its precise molecular mechanism remains unclear, as Retro-2 has also been implicated in perturbing ER export through engagement with Sec16A (*34*). We therefore sought to determine whether Retro compounds share a common mechanism to broadly disrupt TA protein biogenesis.

In this study, we demonstrate that Retro-1, like Retro-2, inhibits TA protein handover from SGTA to Get3. Strikingly, we find that Retro-1, Retro-2, and the optimized analog DHQZ36.1, achieve inhibition by stabilizing rather than disrupting the pretargeting complex. Cryo-electron microscopy (cryo-EM) structures reveal that Retro compounds bind a cryptic, conserved pocket in Get3. Acting as allosteric modulators, the compounds stabilize a Get3 conformation that is nonproductive for TA-protein binding, thereby sequestering the pretargeting complex in a stalled state. These structures fully explain prior structure-activity relationships (SAR) and are corroborated by biochemical analyses. Together, our findings define the mechanistic target of Retro analogs and establish an atomic-level framework for targeting transient complexes through allosteric stabilization.

### Retro compounds target Get3 and stabilize the pretargeting complex

Retro-1, Retro-2, and DHQZ36.1 were synthesized as previously described (Fig. 1B) (*7*, *14*, *15*). To test whether Retro-1 inhibits GET pathway-mediated TA targeting, we used an established dual-fluorescence TA reporter in which GFP is linked via a self-cleaving peptide to RFP fused to the Sec61B transmembrane domain (*27*). In this assay, the RFP:GFP ratio reports TA protein biogenesis efficiency. Genetic deletion of Get3 reduced the RFP:GFP ratio, consistent with destabilization of the fluorescent reporter due to impaired TA-protein insertion. Retro-1 reduced the ratio in a dose-dependent manner, similar to what was previously observed for Retro-2 analogs (Fig. 1C and fig. S2) (*27*). We next assessed TA handover using an *in vitro* crosslinking assay to monitor transfer of the Stx5 transmembrane domain (Stx5_TMD_) from SGTA to Get3 in the presence of Get4, Bag6, and Ubl4a (*27*, *43*). Retro-1 reduced formation of the Get3-Stx5_TMD_ adduct, indicating inhibition of TA-protein transfer from SGTA to Get3 (Fig. 1D and fig. S3).

To determine the molecular basis for Retro compound inhibition of TA handover, we overexpressed and purified the human homolog of Get3 (Get3, residues 22-348) (fig. S4) and performed differential scanning fluorimetry (DSF). ATP increased the melting temperature of Get3, consistent with the expected stabilization of its closed state (fig. S5). The optimized compound DHQZ36.1 enhanced this thermal shift, suggesting its specific binding to Get3 (fig. S5). Retro-1 and Retro-2 did not significantly alter the melting temperature of Get3, perhaps highlighting the potential for optimization of these compounds. We next assessed the effect of the compounds on ATPase activity. Under saturating conditions, the V_max_ for Get3 ATPase activity was 150 ± 70 nmol P_i_/min/µmol (fig. S6), substantially lower than that reported for yeast Get3 (V_max_ = 4,500 ± 500 nmol Pi/min/µmol) (*44*, *45*), suggesting a more tightly regulated ATPase cycle in the human system (*46*). Retro-1, Retro-2, and DHQZ36.1 had no significant effect on Get3 ATPase activity, indicating that these compounds do not inhibit hydrolysis directly but instead act through an alternative mechanism (fig. S6).

We next examined whether Retro analogs influence the interaction between the pretargeting components Get3, Get4, and Bag6 using a purified, coexpressed complex of human Get4 (Get4, residues 23-304) and Bag6 (Bag6, residues 1000-1054) referred to as Get4/Bag6 (*47*). Pretargeting complex formation was measured using capture assays (*48*), in which untagged Get3 was incubated with tagged Get4/Bag6 on beads, and bound Get3 was quantified after washing (Fig. 1E). As previously observed in yeast (*48*), Get3 capture was ATP-dependent (∼0.3 Get3 per Get4/Bag6) (Fig. 1E). Strikingly, addition of Retro-1, Retro-2, or DHQZ36.1 increased Get3 capture to near-stoichiometric levels (∼1 Get3 per Get4/Bag6) (Fig. 1E and fig. S7). Retro compounds stabilize the Get3–Get4/Bag6 interaction without blocking ATP hydrolysis, suggesting that they might function like molecular glues to stabilize a protein-protein interaction (*49*), rather than orthosteric inhibitors. This hypothesis prompted an investigation into the structural basis of this mechanism.

### Cryo-EM structure of Get3–Get4/Bag6 complex bound to Retro-2

We determined the cryo-EM structure of a simplified human pretargeting complex bound to Retro-2. Using the same purified constructs as in the capture assay, Get3 and Get4/Bag6 were incubated with ATP and Retro-2 prior to vitrification. Single-particle analysis yielded a Coulomb potential map refined to an overall resolution of 2.9 Å (Fig. 2A, fig. S8, and table S2). The density revealed a Get3 dimer stoichiometrically associated with Get4/Bag6 in an approximately C2-symmetric assembly (Fig. 2B, S8). C2 symmetry was applied in the final refinement. The quality of the density enabled near-complete modeling of Get3 (α-helices H1-H12 and β-strands B1-B7), Get4 (α-helices h1-h16), and the Bag6 C-terminus, with only residues 208-212 of Get3 remaining unmodeled (Fig. 2B–C and fig. S9).

**Fig. 2.**
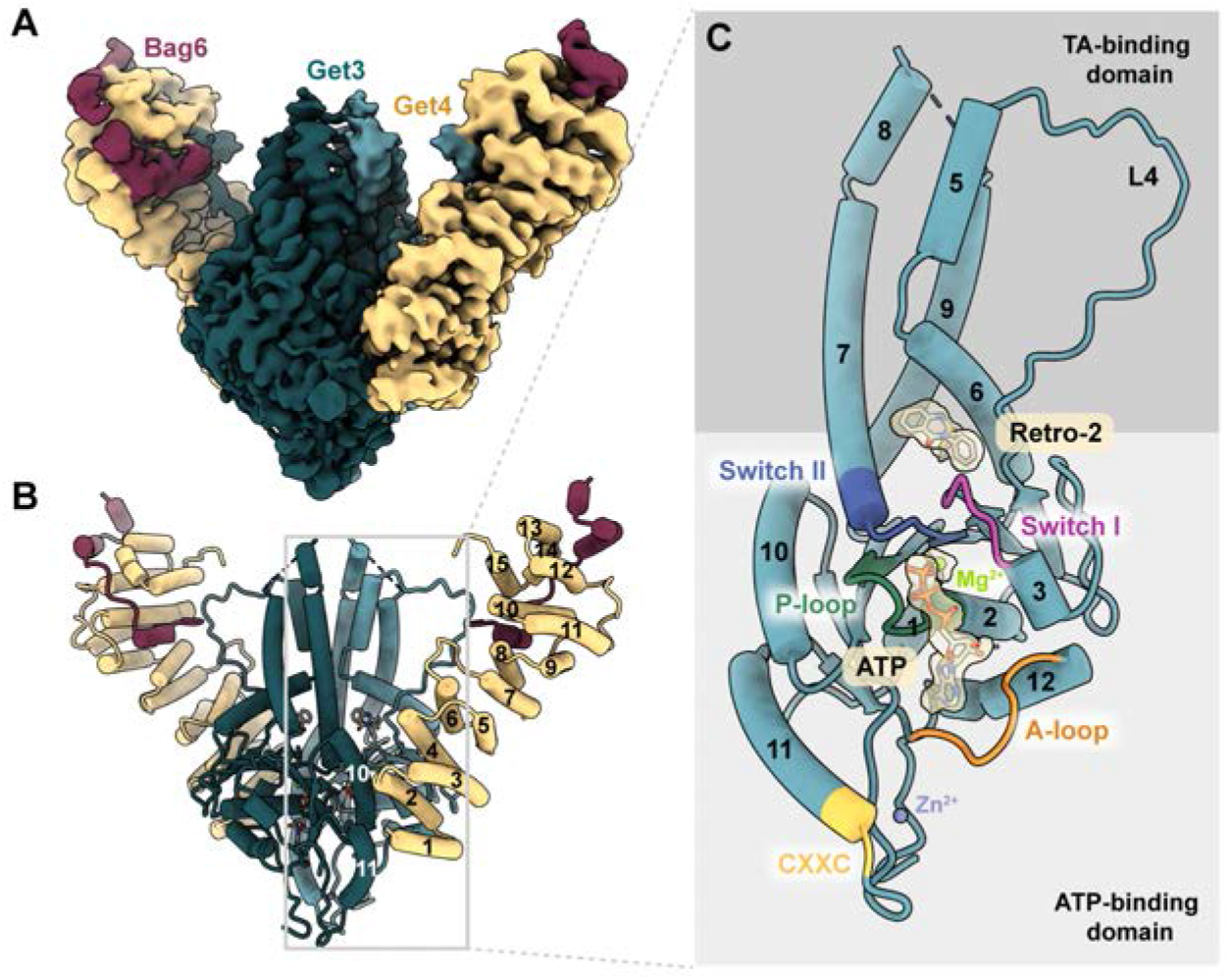
Cryo-EM structure of the Get3–Get4/Bag6 complex bound to Retro-2. (**A**) Cryo-EM density map of the Get3–Get4/Bag6 complex bound to Retro-2, with Get3 shown in teal, Get4 in yellow, and Bag6 in maroon. (**B**) Atomic model of the Get3–Get4/Bag6 simplified pretargeting complex bound to Retro-2, colored as in (A). (**C**) Structure of a single Get3 monomer annotated with helix numbers and conserved Walker A ATPase motifs; cryo-EM density is depicted for Retro-2, ATP, and Mg^2+^.

Get3, consisting of an ATP-binding domain and a TA-binding domain, adopts a closed conformation (Fig. 2B,C). Helices H10 and H11 in Get3 engage Get4 helices h1 and h2 (Fig. 2B and fig. S9), consistent with the previously reported chimeric metazoan (*Danio rerio* Get3 bound to human Get4/Bag6/Ubl4a) and yeast complexes (*38*, *50*). Notably, the flexible L4 loop of Get3 (residues 104-126) is fully resolved and forms a secondary interaction interface with Get4 helices h10 and h15 and the Bag6 C-terminus through electrostatic and hydrophobic contacts (Fig. 2B,C, and fig. S9) (*38*).

After modeling protein, ATP, and metal cofactors (Mg^2+^, Zn^2+^), a trilobed density remained at the interface between the ATP- and TA-binding domains of each Get3 monomer (Fig. 2C) While a racemic mixture of Retro-2 was added to the sample, only the cyclized (*S*)-enantiomer of Retro-2 fit the density, revealing the preferred stereochemistry of the active compound (Fig. 2C, 3A) (*13*). Retro-2 occupies a hydrophobic cryptic pocket formed primarily by helices H6 and H7 and is stabilized by hydrogen bonds with Ser147 and His172, extensive hydrophobic contacts, and a T-shaped aromatic interaction with Phe179 (Fig. 3A-D) (*51*). Located at the interface of the ATP- and TA-binding pockets, this cryptic pocket defines a previously unrecognized allosteric site in Get3, hereafter referred to as the Retro-binding site.

**Fig. 3.**
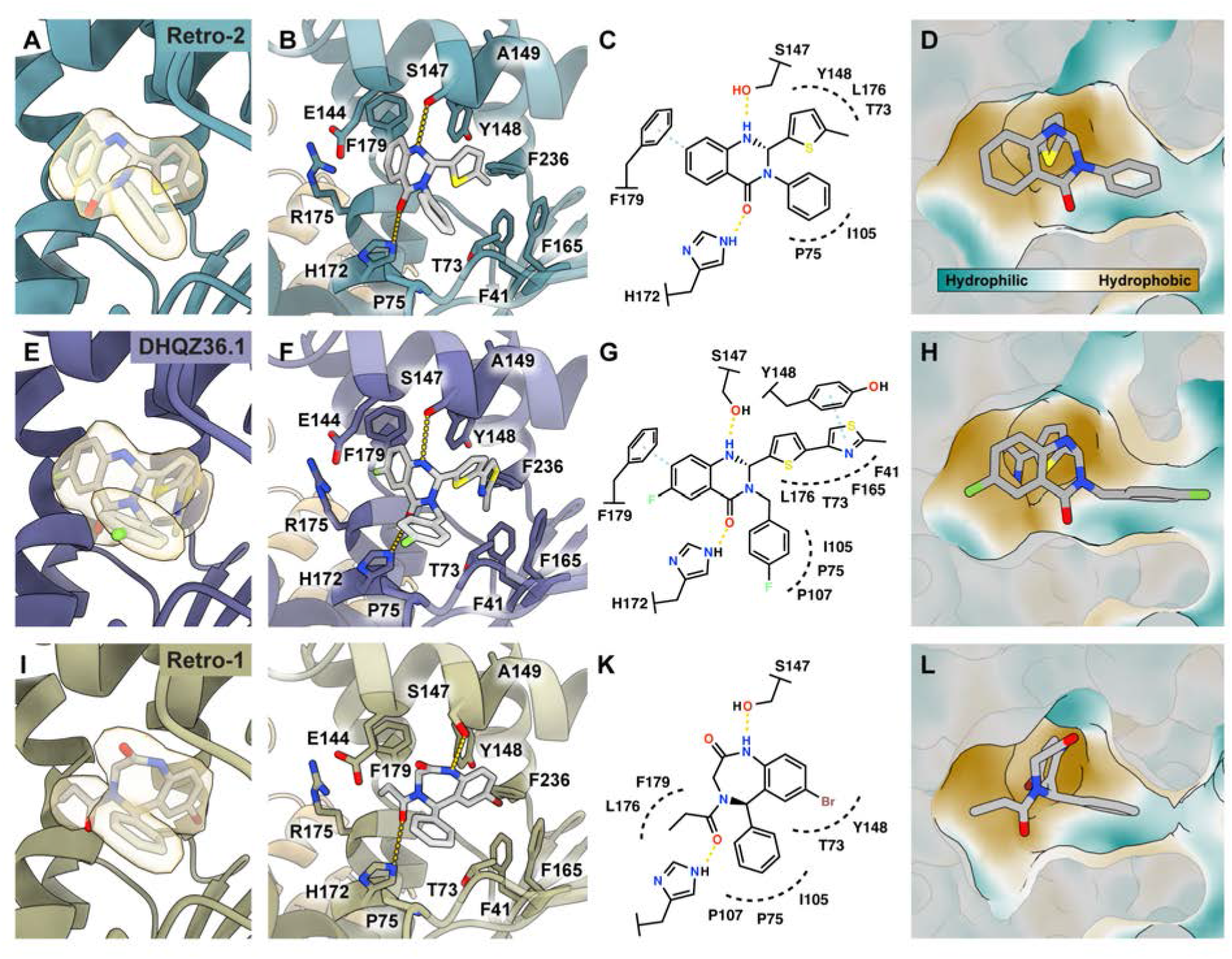
Retro-1 and DHQZ36.1 bind a cryptic Get3 pocket. (**A**,**E**,**I**) Cryo-EM density with corresponding models for (A) Retro-2, (E) DHQZ36.1, and (I) Retro-1. (**B**,**F**,**J**) Close-up views of the Retro-binding pocket with key pocket residues labeled and hydrogen bonds indicated. (**C**,**G**,**K**) Schematic representations of the Retro-binding pocket for each compound. (**D**,**H**,**L**) Surface representations of the Retro compound binding pocket colored by hydrophobicity (teal, hydrophilic; gold, hydrophobic).

### Retro-1, Retro-2, and DHQZ36.1 bind a cryptic Get3 pocket

Following incubation with the optimized Retro-2 derivative DHQZ36.1 or with Retro-1, the Get3–Get4/Bag6 complex was vitrified under the same conditions as the Retro-2 complex, yielding cryo-EM structures at 2.8 and 2.7 Å resolution, respectively (fig. S10–11 and table S2). In both cases, Get3 and Get4/Bag6 adopt conformations identical to the Retro-2-bound complex. Trilobed density at the same cryptic pocket (Fig. 3E,I and fig. S11) confirmed that DHQZ36.1 and Retro-1 target the same Retro-binding site in Get3, despite having distinct chemical structures (Fig. 3E-L, Fig. 4A, and fig. S12, S16).

**Fig. 4.**
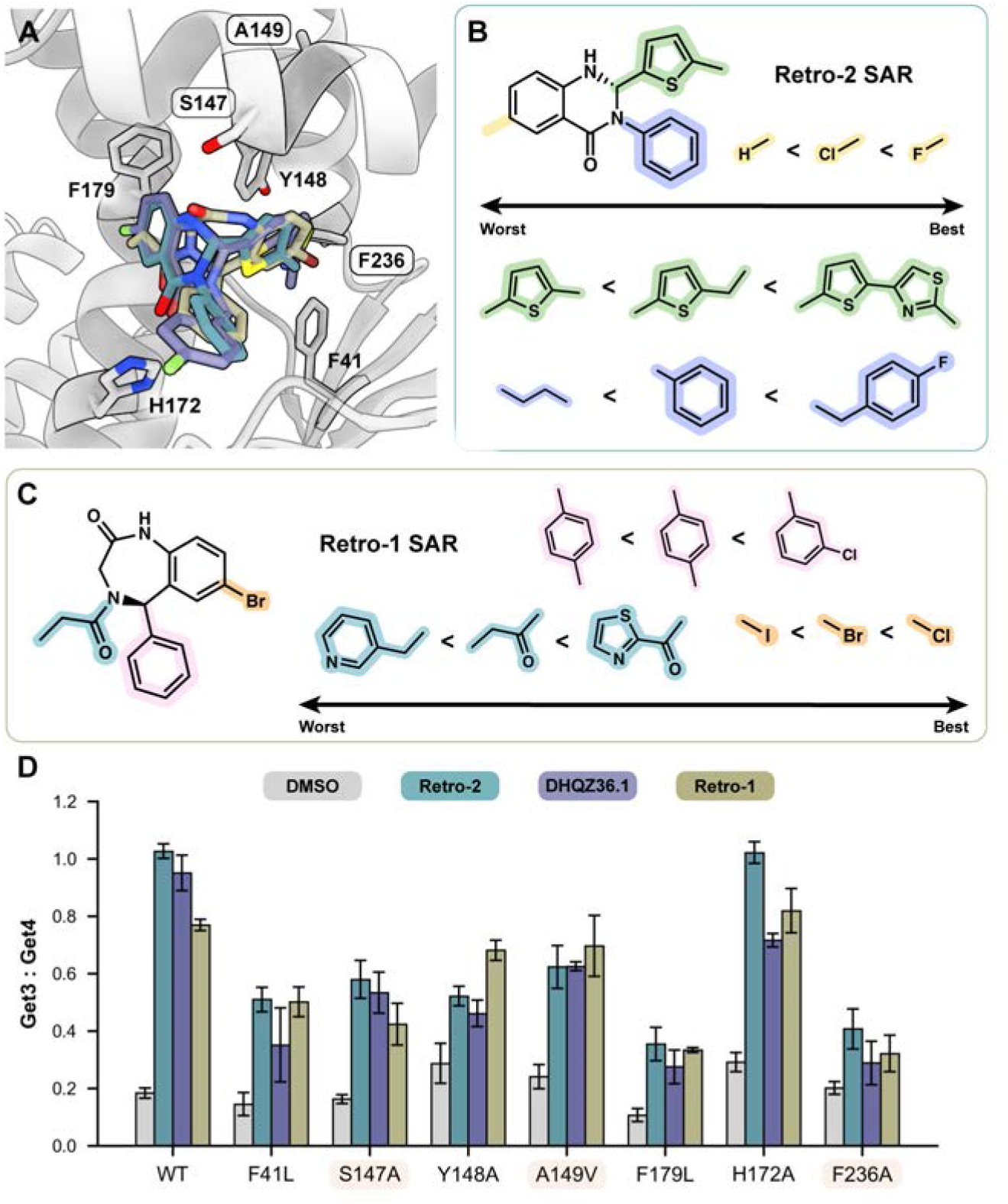
Mutational analysis of residues involved in Retro compound binding. (**A**) Overlay of Retro compounds in the Retro-2 binding pocket (grey): Retro-1 (green), Retro-2 (teal), DHQZ36.1 (purple). Key interacting residues are labeled, with compound-resistant mutants boxed. (**B**) Representative SAR trends for Retro-2, full SAR trends are shown in fig. S13. (**C**) Representative SAR trends for Retro-1; full SAR trends are shown in fig. S15. (**D**) Capture assays showing Get3–Get4/Bag6 complex formation for Get3 mutants in the presence of ATP and the indicated compounds. Data are shown as mean ± SE (≥3 replicates per condition).

For both Retro-2 and DHQZ36.1, the (*S*)-enantiomer engages the same core hydrogen-bonding network with Ser147 and His172 and makes a T-shaped aromatic interaction with Phe179 (Fig. 3F–G). The thiazole moiety of DHQZ36.1 additionally forms hydrophobic contacts with Phe41, Phe165, and Phe236, as well as a face-to-face aromatic interaction with Tyr148 (Fig. F–G). This explains prior SAR observations: thiophene C2 substitution enhances potency (*e.g*., ethyl substitution increases hydrophobic contacts) and bulky thiazole substituents more fully occupy the pocket and engage Tyr148 (Fig. 3H, 4B and fig. S13) (*13–15*, *52*). Likewise, the increased potency of DHQZ36.1 against *Leishmania amazonensis* infection is consistent with additional contacts made by the 4-fluorobenzyl moiety relative to the smaller phenyl group of Retro-2 (Fig. 3H, 4B and fig. S13). Although optimization of DHQZ36.1 was structure-blind, the cryo-EM structure rationalizes its superior potency and also explains why other molecules optimized from the quinazolinone scaffold (*e.g.*, Retro-2.1) similarly exhibit enhanced potency (fig. S13) (*13*, *14*).

The (*R*)-enantiomer of Retro-1 was fit into the density and forms a hydrogen-bonding network with Ser147 and His172 similar to that observed for Retro-2 and DHQZ36.1 (Fig. 3J,K). The ligand conformation is clearly resolved in the density (Fig. 3I) and corroborates DSF measurements in which (*R*)-Retro-1 exhibited a larger thermal shift than the (*S*)-enantiomer, consistent with higher affinity to Get3 (fig. S15). Retro-1 maintains hydrophobic contacts with Phe179, Pro75, and Ile105 (Fig. 3J,K). Its phenyl ring extends into the hydrophilic region of the pocket (Fig. 3L), explaining SAR trends: *meta*-chloro substitution improves potency while *para*-methyl substitution reduces it (Fig. 4C and fig. S15) (*11*, *22*). Replacement of the propionyl moiety of Retro-1 with an *N*-thiazole-2-carbonyl substituent further enhances potency, suggesting opportunities for structure-guided optimization to better complement pocket architecture (Fig. 4C and fig. S15) (*22*). Likewise, for the benzo moiety of Retro-1, the observed correlation between potency and halogen radius (Cl > Br > I) indicates that steric complementarity within the pocket is a stronger determinant of activity than electronic effects (Fig. 4C and fig. S15).

### Key hydrophobic Get3 residues mediate Retro compound activity

In identifying Get3 as a potential Retro target (*27*), CRISPR-X screens against DHQZ36.1 yielded a range of compound-resistant mutants, including S147R, A149V, and F236L (Fig. 4A and fig. S17). Ser147 and Phe236 directly contribute to the Retro-binding pocket, with Ser147 forming a hydrogen bond to the carbonyl. Ala149 is positioned above H6 relative to the binding pocket, adjacent to residues Ser147 and Tyr148, suggesting it affects conformational stability. Additional mutations occur distal to the Retro-binding pocket, connecting helix H7 to the ATP-binding site (fig. S17).

To define the role of binding pocket residues in Retro-dependent stabilization of the pretargeting complex, we generated point mutants and measured Get3 capture by Get4/Bag6 in the presence of Retro-1, Retro-2, and DHQZ36.1. Mutations that reduced compound binding were expected to disrupt Retro-enhanced capture without affecting the basal Get3-Get4/Bag6 interactions. Most mutants retained ATP-dependent capture similar to that of the wild type (Fig. 4D, S18). Those lacking measurable ATP-dependent capture were excluded from further analysis. To control for modest differences in basal capture among the remaining mutants, fold changes in the Get3:Get4/Bag6 ratio were calculated relative to basal ATP-dependent capture, with statistical significance relative to wild-type capture reported as −log_10_(*p*-values) (fig. S18).

Consistent with their identification in the CRISPR-X screen and their positioning at or near the Retro-binding pocket, mutations at Ser147, Ala149, and Phe236 reduced Retro-enhanced Get3 pulldown (Fig. 4D and fig. S18). Mutation of Tyr148, which makes an aromatic interaction with the DHQZ36.1 thiazole, similarly diminished capture (Fig. 4D and fig. S18). Substitution of hydrophobic residues Phe41 and Phe179 with leucine also attenuated complex formation across all three compounds, underscoring their central role in compound binding (Fig. 4D and fig. S18). Alanine substitution of His172, which forms a hydrogen bond, produced only a modest reduction in capture (Fig. 4D and fig. S18). Combined, Retro-dependent pretargeting complex stabilization is primarily mediated by hydrophobic and shape-complementary dispersion interactions.

### Retro compounds stabilize a nonproductive Get3 conformation to prevent TA handover

Get4/Bag6 binding to Get3 facilitates conformational changes that stabilize H6 in a downward position, promoting rearrangement of the remaining TA-binding helices into the hydrophobic TA-binding groove (Fig. 5A) (*37*, *38*). Although Get4 engages Get3 in a manner consistent with previous structures, Retro compound binding at the interface between the ATP- and TA-binding domains prevents H6 from descending, trapping Get3 in an H6-up conformation rather than the expected H6-down state associated with the pretargeting complex (Fig. 5B and fig. S19). This conformation allows the flexible Get3 L4 loop to form a rigid interaction with Get4, contributing to stabilization of the complex. In this stalled configuration, the hydrophobic TA-binding groove cannot form. The resulting architecture resembles the helix organization observed in the ATP-bound *Giardia intestinalis* Get3 structure (Fig. 5B and fig. S20) (*37*).

**Fig. 5.**
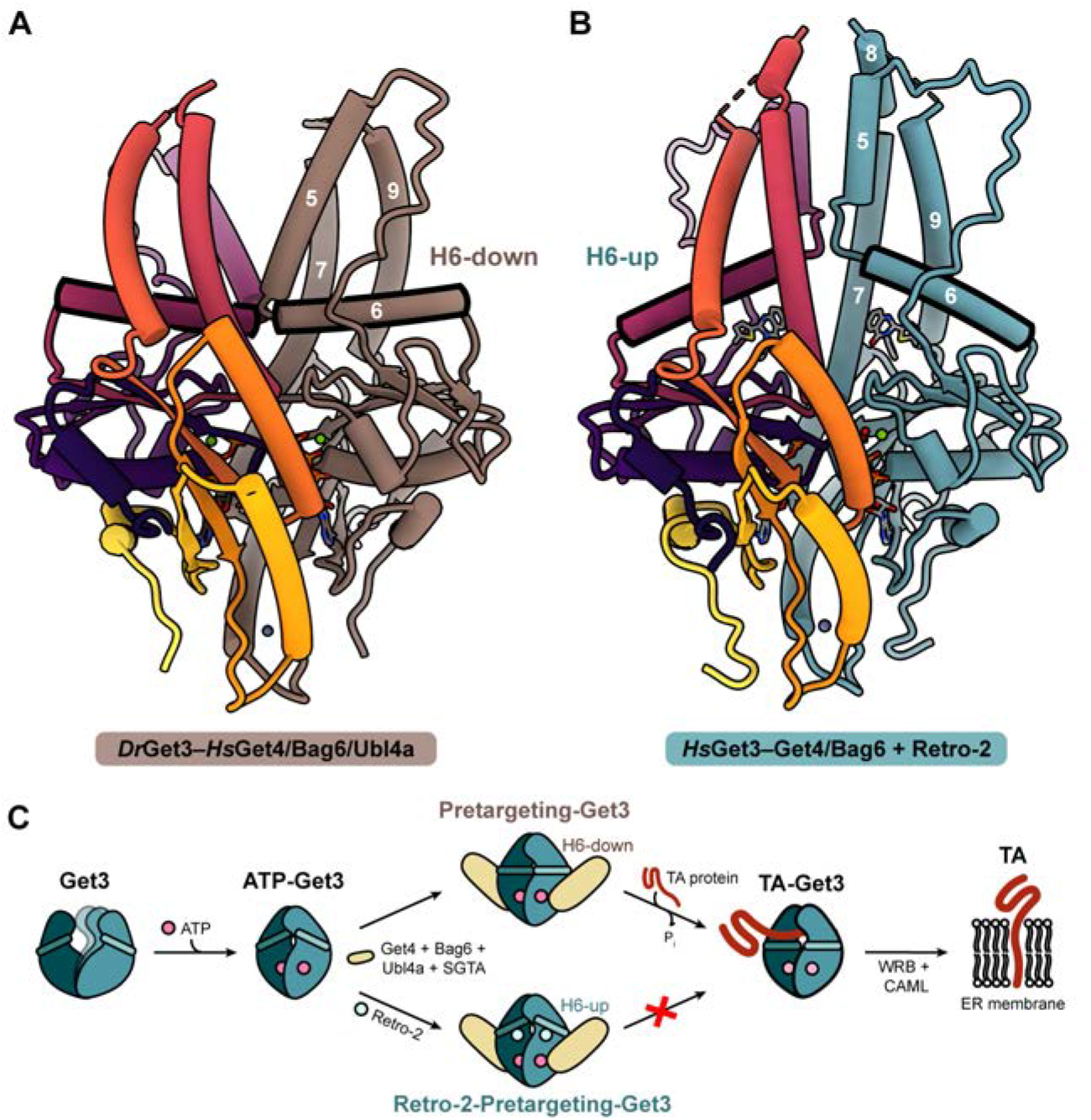
Retro-2 stabilize a nonproductive Get3 conformation to prevent handover. (**A**) Structure of the chimeric pretargeting complex (PDB ID: 7ru9) composed of zebrafish Get3 and human Get4/Bag6/Ubl4a. Only Get3 is shown, with one monomer in brown and the other colored sequentially using the inferno palette (N-terminus, purple; C-terminus, yellow). Helix 6 adopts a downward conformation. Zebrafish and human Get3 share 99% sequence similarity, making this structure a suitable reference. (**B**) Retro-2-bound Get3–Get4/Bag6 complex. Only Get3 is shown, with one monomer in teal, the other colored sequentially using the inferno palette. Helix 6 adopts an upward conformation. (**C**) Model of Get3 conformational transitions during ATP binding: ATP stabilizes a closed conformation, with formation of the pretargeting complex resulting in helix 6 moving down (top), and TA loading driving nucleotide hydrolysis and release. Retro-2 stabilizes the pretargeting complex (bottom), stalling Get3 and blocking TA transfer to Get3.

Retro compound binding alters the Get3 dimer interface. The hydrogen bond with His172 disrupts the observed interdimer histidine stacking and creates new interactions between Arg175 and Glu144, as well as between Pro140 of opposite monomers (fig. S19) (*38*, *48*, *50*). By simultaneously remodeling dimer contacts, occluding the hydrophobic groove, and stalling H6 movement, Retro compounds trap the pretargeting complex in a handover-incompetent state, thereby blocking TA protein biogenesis.

### Allosteric stalling of the GET pathway by Retro compounds

Retro compounds bind at a critical communication point between nucleotide recognition and TA protein capture, trapping Get3 in a nonproductive complex with other components of the GET pathway. In this respect, they resemble molecular glues, which stabilize protein-protein interactions (*53–55*). Unlike molecular glues, however, Retro compounds function as allosteric modulators, stabilizing transient interactions through binding at a site distinct from the protein-protein interface (*56*, *57*).

Functionally, trapping a handoff step in the GET pathway is sufficient to block TA protein biogenesis, including SNAREs required for retrograde transport (*27*, *30*, *34*). Retro compounds act on their targets indirectly, perturbing host trafficking by modulating pathway dynamics. These findings highlight transient complexes in the secretory pathway as an underexplored class of therapeutic targets. Related strategies have shown promise by targeting other central nodes in secretion, including the Sec61 translocon, DPAGT1-mediated N-linked glycosylation, and ARF1-dependent Golgi transport (*58–60*).

From a therapeutic perspective, Retro compounds have previously been reported to be well tolerated in cells and mice, suggesting that disruption of the GET pathway may be pharmacologically tractable (*10*). Our structural analysis of both the original hits and the optimized analog DHQZ36.1 suggests that binding is primarily driven by shape complementarity and van der Waals interactions, providing a framework for structure-guided optimization. More broadly, allosteric stabilization of transient interactions may offer advantages over orthosteric inhibition, including increased specificity and tunability (*61*).

Two decades after their discovery in the same screen, our findings uncover a common mechanism for two structurally distinct classes of Retro compounds: allosteric stabilization of a nonproductive Get3 state that blocks TA protein biogenesis. Beyond defining the mechanism of Retro compounds, this work demonstrates that transient states within cellular pathways can be selectively targeted by small molecules, providing a framework for pharmacologically modulating dynamic cellular pathways.

## Acknowledgments

1. J. Sello is an investigator of the Chan Zuckerberg Biohub San Francisco. We are grateful to S. Mahajan, A. Barlow, M. Fry, C. Wells, B. Y. Kaudeer, J. Kirsh, G. Baron, R. Voorhees, T.-F. Chou, P. Bjorkman, S.-O. Shan, R. Thompson, S. Shoemaker, S. Marqusee, and the entire Clemons-Rees group for lively discussion and advice. Cryo-EM was performed in the Beckman Institute Resource Center for Transmission Electron Microscopy at Caltech, and we are indebted to S. Chen, T. Brittain, and W. Floriano for microscope and computing expertise. We are also grateful to R. Sarpong for assistance with enantiomeric separation. Finally, we thank L. Piro and T. Safar for technical assistance, as well as Caltech administrative and custodial staff for their essential support.

## Author contributions

Conceptualization: JAL, WMC

Methodology: JAL, XG, CJC, VSR, VGR, YEL, VD, JKS, WMC

Investigation: JAL, XG, CJC, VSR, VGR, YEL, AHN, VD, JKS, WMC

Visualization: JAL, CJC, VSR, WMC

Funding acquisition: WMC

Project administration: JKS, WMC

Supervision: JKS, WMC

Writing – original draft: JAL, WMC

Writing – review & editing: JAL, XG, CJC, VSR, VGR, YEL, AHN, VD, JKS, WMC

## Data and materials availability

All experimental data are available in the main text or the supplementary materials. Coordinates and their experimental maps have been deposited to the RCSB and EMDB, with accession numbers XXXX and EMD-XXXXX.

## Materials and Methods

### Retro compound synthesis

Retro-1, Retro-2, and DHQZ36.1 were synthesized as previously described (*11*, *14*, *15*). *Retro-1*: ^1^H NMR (400 MHz, DMSO) δ ppm: 10.08 (d, 1H), 7.83 (dd, 1H), 7.54 (td, 1H), 7.42 – 7.18 (m, 3H), 7.10 – 6.93 (m, 3H), 6.58 (d, 1H), 4.32 – 3.91 (m, 2H), 2.66 – 2.53 (m, 1H), 2.47 – 2.30 (m, 1H), 1.01 (dt, 3H). *Retro-2:* ^1^H NMR (400 MHz, CDCl_3_) δ ppm: 7.73 (d, 1H), 7.61 (d, 1H), 7.40-7.32 (m, 2H), 7.32 (m, 3H), 7.32–7.30 (m, 1H), 6.83-6.79 (d, 1H), 6.78 (t, 1H), 6.77 (d, 1H), 6.73 (d, 1H), 6.55 (d, 1H), 2.31 (s, 3H). *DHQZ36.1:* ^1^H NMR (400 MHz, CDCl_3_) δ ppm: 7.71 (dd, 1H), 7.33–7.24 (m, 3H), 7.21 (d, 1H), 7.15 (s, 1H), 7.08–6.98 (m, 3H), 6.85 (d, 1H), 6.60 (dd, 1H), 5.78 (s, 1H), 5.57 (d, 1H), 3.88 (d, 1H), 2.73 (s, 3H). All spectroscopic data are consistent with those reported in the literature (*11*, *14*, *15*). Retro-1 enantiomers were separated using a Lux 5 µm i-Amylose-3 LC 100 x 10 mm column

### Retro compound enantiomer separation

An HPLC system consisting of a Shimadzu CBM-40 controller, LC-20AR pumps, and an SPD-M40 UV detector was used for enantiomeric separation studies. Separation of the Retro-1 enantiomers was performed using a Lux® 5 µm i-Amylose-3 column (100 x 10 mm). The eluent consisted of a gradient of water and acetonitrile, delivered at a flow rate of 5.0 mL/min. Samples were injected at a volume of 30 µL, and chromatograms were recorded at 222 nm.

### TA reporter assay

Previously, a doxycycline-inducible cassette was engineered to express GFP linked by a self-cleaving P2A peptide to a RFP fused to a C-terminal Sec61B transmembrane domain (TMD) sequence (GFP-2A-RFP-Sec61_TMD_) (Morgens 2019). Wild-type (WT) and Get3^KO^ HEK293T cell lines with the GFP-2A-RFP-Sec61B_TMD_ were pre-treated with the indicated compounds for 1 hour prior to induction with doxycycline for approximately 18 h. Fluorescence activated cell sorting (FACS) was performed to determine RFP:GFP ratio.

### TA handover assay

A ^35^S-methionine labeled model tail-anchored substrate consisting of the cytosolic domain of Sec61B and the TMD of Stx5 followed by C-terminal opsin tag (STX5_TMD_) was translated in the presence of recombinant SGTA (final concentration of ∼14 µM) in a PURE translation system (*39, 56*). Translation was carried out at 37 °C for 90 min. SGTA bound to Stx5_TMD_ (SGTA∼Stx5_TMD_) was purified by size-separation through a 5-25% sucrose gradient. SGTA∼Stx5TMD complex was then incubated on ice with zebrafish Get3 and human Ubl4a, Bag6 (1032-1132), and Get4 in the presence of MgATP and 20 µM Retro-1 to monitor transfer of Stx5_TMD_ from SGTA to Get3. Completed reactions were warmed at 32 °C for 1 min and subjected to chemical crosslinking (XL) with 0.250 mM bismaleimidohexane (BMH) on ice for 1 h. Crosslinked samples were resolved by SDS-PAGE and visualized by autoradiography and Coomassie Blue staining.

### Cloning and expression of *Hs*Get3

For biochemical and structural studies, a truncated version of the *Homo sapiens* Get3 gene (residues 22-348) was cloned into a pET33b vector with N-terminal hexahistidine and SUMO fusion protein tags. A full-length *Hs*Get3 construct (residues 1-348) was cloned in the same manner and used to confirm that truncation did not affect catalytic activity. Point mutations were introduced into the *Hs*Get3 construct using Q5 site-directed mutagenesis (New England Biolabs). Wild-type and mutant *Hs*Get3 were expressed in *E. coli* NiCo21(DE3) cells (New England Biolabs) which were grown in baffled flasks containing 2xYT media (16 g/L tryptone, 10 g/L yeast extract, 5 g/L NaCl) supplemented with kanamycin (50 μg/mL) at 37 °C with shaking at 250 rpm. Protein expression was induced at an optical density (OD_600_) of ∼0.6 with 0.7 mM isopropyl-β-d-thiogalactopyranoside (IPTG), after which the temperature was reduced to 20 °C. Cells were harvested after 18 h by centrifugation in a JLA-8.1 rotor at (9,000 x g) for 10 min.

### Purification of *Hs*Get3

Cell pellets were resuspended in 10 volumes of Get3 lysis buffer (50 mM Tris pH 7.5, 300 mM NaCl, 20 mM imidazole, 10 mM β-mercaptoethanol, 1 mM PMSF, 1 mM benzamidine) and lysed using an M-110L microfluidizer (Microfluidics) with three passes at ∼17,500 psi. The lysate was clarified by ultracentrifugation at 38,000 x g in a Type 45 Ti rotor for 45 min at 4 °C and subsequently incubated with 2 mL of nickel-nitrilotriacetic acid (Ni-NTA) agarose (Qiagen) for 2 h at 4 °C with gentle rocking. Resin was washed with 100 mL of Get3 lysis buffer and His-tagged Get3 protein was eluted with 24 mL Get3 elution buffer (50 mM Tris pH 7.5, 150 mM NaCl, 300 mM imidazole, 10 mM β-mercaptoethanol, 1 mM PMSF, 1 mM benzamidine). Affinity tags were removed by overnight digestion with ULP1 protease in 3.5 kDa MWCO SnakeSkin dialysis tubing (Thermo Fisher Scientific) in Get3 dialysis buffer (20 mM Tris pH 7.5, 150 mM NaCl, 10 mM β-mercaptoethanol) at 4 °C. The dialyzed sample was passed over Ni-NTA resin to remove the cleaved His-SUMO tag. The untagged protein was concentrated to ∼500 μL using an Amicon Ultra centrifugal filter (30 kDa MWCO, 2,500 x g) and further purified by size-exclusion chromatography on a Superdex 200 Increase 10/300 GL column (Cytiva) in Get3 sizing buffer (20 mM Tris pH 7.5, 75 mM NaCl, 10 mM β-mercaptoethanol). Fractions were analyzed by SDS-PAGE and pure fractions were pooled, concentrated, and either used immediately or flash-frozen.

### Differential Scanning Fluorimetry

All DSF experiments used 8-well TempAssure, Optical Cap PCR tubes. BIODIPY FL L-cysteine (Invitrogen) was diluted in DMSO and was at 4 μM in the final reaction. All small molecules were also diluted in DMSO and were at 8 μM in the final reaction. The final reaction volume for all reactions contained 5% DMSO, including the small-molecule and dye additions. An ATP mixture was prepared by mixing an equimolar solution of ATP salt and MgCl_2_ in the reaction buffer. To ensure that ATP was in excess throughout the experiments, a final ATP/Mg^2+^ concentration of 16 μM was used for all indicated reactions. The final concentration of HsGet3 was 4 μM. A BioRad C100 Touch thermal Cycler^TM^ with a CFX96 Real-Time System^TM^ was used for all measurements. Measurements started with a 15-minute hold at 4 °C, after which the temperature was increased at 1 °C per minute, with a measurement after each minute until 95 °C is reached. The data was analyzed in Microsoft Excel and TPPU_DSF public server from the Macias Lab (*62*). The melting temperature was determined using the Boltzmann derivative, and the significance was calculated using a t-test.

### Cloning and expression of *Hs*Get4/Bag6

Truncated versions of the *Hs*Get4 (23-305) and *Hs*Bag6 (1006-1059) genes were inserted into the multiple cloning site of a pACYC vector, with an N-terminal hexahistidine tag on Bag6. The plasmid was expressed in *E. coli* NiCo21(DE3) cells which were grown in baffled flasks containing 2xYT media supplemented with chloramphenicol (35 μg/mL) at 37 °C with shaking at 250 rpm. Expression was induced at an OD_600_ of ∼0.6 using IPTG, and cells were harvested after 3 h by centrifugation in a JLA-8.1 rotor at (9,000 x g) for 10 min.

### Purification of *Hs*Get4/Bag6 complex

Cell pellets were resuspended in 10 volumes of Get4/Bag6 lysis buffer (20 mM Mops pH 7.2, 300 mM K⋅glutamate, 5 mM MgOAc, 20 mM imidazole, 5 mM β-mercaptoethanol) and lysed using a M-110L microfluidizer with three passes at ∼17,500 psi. The lysate was clarified by ultracentrifugation at 38,000 x g in a Type 45 Ti for 45 min at 4 °C and subsequently incubated with 2 mL of Ni-NTA agarose for 1 h at 4 °C with gentle rocking. Resin was washed with 100 mL of Get4/Bag6 lysis buffer and the His-tagged Get4/Bag6 complex was eluted with 12 mL Get4/Bag6 elution buffer (20 mM Mops pH 7.2, 150 mM K⋅glutamate, 300 mM Imidazole, 5 mM β-mercaptoethanol). For structural studies, the affinity tag was then removed by overnight digestion with TEV protease in 3.5 kDa MWCO SnakeSkin dialysis tubing in Get4/Bag6 dialysis buffer (20 mM Mops pH 7.2, 100 mM K⋅glutamate, and 10 mM β-mercaptoethanol) at room temperature. This step was omitted for complexes prepared for capture assays. The sample was concentrated to ∼500 μL using an Amicon Ultra centrifugal filter (10 kDa MWCO, 2,500 x g) and purified by size-exclusion chromatography on a Superdex 75 Increase 10/300 GL column (Cytiva). Fractions were analyzed by SDS-PAGE and pure fractions were pooled, concentrated, and either used immediately or flash-frozen.

### Get3 ATPase activity assay

ATPase assays were performed using the Malachite Green Phosphate Assay Kit (Sigma-Aldrich), following a protocol adapted from Rule *et al*. (2016). Reactions (500 µL) were carried out in assay buffer (200 mM HEPES pH 7.5, 150 mM NaCl, 5 mM β-mercaptoethanol) containing 20 µM Get3, 5 mM MgCl_2_, and 5 mM ATP. DMSO, Retro-1, Retro-2, or DHQZ36.1 were included at 500 µM. Reactions were incubated at 37 °C for 1 h, with aliquots taken prior to reaction initiation and at 15 min intervals thereafter. At each timepoint, 10 µL of the reaction was diluted into 140 µL of buffer on ice and immediately quenched by flash-freezing in liquid nitrogen. Samples were transferred to −80 °C for at least 15 min to ensure complete freezing prior to phosphate quantitation. Frozen samples were thawed and transferred to a 96-well clear-bottom plate, alongside phosphate standards. Sample and standard volumes were 50 µL, and measurements were performed in technical duplicate. Malachite green-molybdate working reagent (20 µL) was added to each well and mixed. After incubation for 30 min at room temperature, absorbance was measured at 620 nm using an Infinite M Nano plate reader (Tecan).

### Capture assay

Capture assays were performed following a protocol adapted from Gristick *et al.* (2015). His_6_-tagged Get4/Bag6 (1 nmol) was incubated with 10 µL Ni-NTA agarose resin for 1 h at 4 °C in 500 µL binding buffer (20 mM HEPES pH 7.5, 150 mM potassium acetate, 10 mM magnesium acetate, 10% (v/v) glycerol, 25 mM imidazole) and 1 mM ATP where indicated. Get3 (2 nmol) was then added, and the mixture was incubated for an additional 1 h at 4 °C. Following incubation, the reaction was centrifuged at 1,000 x g for 30 s, and the supernatant was removed. The resin was washed by resuspension in 500 µL binding buffer, followed by centrifugation at 1,000 x g for 30 s. This wash step was repeated twice. After the final wash, samples were incubated with 20 µL elution buffer (30 μL of 20 mM Tris, pH 7.5, 100 mM NaCl, 5 mM β-mercaptoethanol and 300 mM imidazole) for 15 min at 4 °C. Bound proteins were collected by centrifugation at 1,000 x g for 30 s. Eluted samples (20 µL) were mixed with 4 µL 6xSDS-PAGE loading dye, resolved on SDS-PAGE gels, and stained with Coomassie Brilliant Blue G-250. Bands were quantified using Image Lab (Bio-Rad).

### Cryo-EM sample preparation and data collection

Drug-bound Get3 complex was prepared by incubating Get3 (60 µM) and Get4/Bag6 (40 µM) in a total volume of 100 µL containing 20 mM HEPES pH 7.5, 150 mM potassium acetate, 10 mM magnesium acetate, 5 β-mercaptoethanol, and 2 mM ATP. Retro-2, DHQZ36.1, or Retro-1 were added to a final concentration of 1 mM, and samples were incubated for 1 h at room temperature. Insoluble drug was removed by centrifugation at 17,000 x g for 5 min. Samples were diluted to a final protein concentration of ∼0.6 mg/mL and applied to glow-discharged holey carbon grids (Quantifoil 1.2/1.3, 300 gold mesh). Grids were glow discharged for 90 s in air at 20 mA using a Pelco easiGlow, Emeritech K100X. 3 µL of sample were applied to grids, which were blotted using a Vitrobot Mark IV (Thermo Fisher Scientific) at 4 °C, 100% humidity for 3.5 s with a blot force of +10, then immediately plunge frozen in liquid ethane cooled by liquid nitrogen. Grids were stored in liquid nitrogen until data collection. Cryo-EM data were collected using SerialEM on a Titan Krios transmission electron microscope (Thermo Fisher Scientific) operating at 300 keV and equipped with a 20 eV energy filter and a Gatan K3 6k x 4k direct electron detector. Movies were collected at a magnification of 130,000x over a defocus range of −0.5 to −2.5 μm, with a total electron flux of 70 e^-^/Å^2^ using correlated double sampling (CDS) mode. For the Retro-2-bound complex, 5,580 movies were collected with 40 frames per movie; 5,498 movies were collected for DHQZ36.1 and 4,777 movies were collected for Retro-1. Data were collected using super resolution mode with a calibrated pixel size of 0.325 Å per pixel. Data collection and processing parameters are summarized in table S2.

### Cryo-EM data processing

Image processing was performed in cryoSPARC (v4.7.1) (*63*) using a similar workflow for all datasets, summarized in Figs. S9-S11. Movies were motion corrected with 2x Fourier cropping, and contrast transfer function (CTF) parameters were estimating using patch CTF estimation. Micrographs were curated based on defocus, CTF fit (<4 Å), ice thickness, and motion statistics. Initial particle picking was performed using blob picking with a minimum particle diameter of 80 Å and a maximum particle diameter of 160 Å. Particles were extracted with a box size of 512 pixels and binned by a factor of 2. Iterative 2D classification was performed using 100 classes, a circular mask diameter of 180 Å, 40 iterations, a batch size of 400 per class, and marginalization over poses and shifts. After strict 2D classification, particles from high-quality 2D classes were used to train a Topaz model for neural network-based particle picking (*64*), and additional good particles were extracted. Ab-initio reconstruction was performed using four classes with an initial resolution of 9 Å, final resolution of 6 Å, initial minibatch size of 400, and final minibatch size of 1000, followed by iterative heterogeneous refinement. 3Particles were reextracted without binning using a box size of 512 pixels. Non-uniform refinement was performed with an initial lowpass filter of 12 Å, two final refinement passes, and minimization over per-particle scale factors. Reference-based motion correction and global and local CTF refinement were performed using a global mask. 3D classification was applied to the semi-final set of particles in order to isolate the major conformation of the complex. As C2 symmetry was observed, the final refinement was performed with C2 symmetry. The overall resolution was estimated using the gold-standard Fourier shell correlation (GSFSC) curve at a cutoff of 0.143.

### Model building

The initial model for the Retro-2 bound complex was assembled from previously reported structures (PDB IDs 4wwr and 7ru9) (*38*, *47*) and AlphaFold3 (6*5*) predictions and docked into the cryo-EM map using rigid body fitting in UCSF ChimeraX (*66*). Model refinement was performed using *phenix.real_space_refine* (*67*), with iterative manual model building and adjustment performed in Coot (*68*). CIF restraints for Retro-2 were generated using Grade2 Web Server (*69*), and the ligand was manually fit into the remaining nonproteinogenic density in the map, followed by model refinement in Coot and Phenix. Unsharpened and sharpened maps were used for model building and interpretation, and the sharpened map was used for refinement.. The TA-binding domain, C-terminus of Get4, and Bag6 exhibited lower resolution and required lower map contour thresholds in order to build in the model. While sidechains are still included in these parts model, their positions could not be definitively assigned. Models for the Retro-1-and DHQZ36.1-bound complexes were built using the refined Retro-2 model as an initial template. Ligand restraints were similarly generated using Grade2, and refinement and manual adjustments were performed in the same manner in Phenix and Coot The final models were validated in Molprobity (*70*). UCSF ChimeraX v.1.9 was used for map and model visualization and figure generation (*66*).

## Data availability

Atomic coordinates of the Get3–Get4/Bag6 complex bound to Retro-2, DHQZ36.1, and Retro-1 are deposited at the Protein Data Bank (PDB) with accession codes XXX. Cryo-EM maps of the Get3–Get4/Bag6 complex bound to Retro-2, DHQZ36.1, and Retro-1 are deposited at the Electron Microscopy Data Bank (EMDB) with accession codes EMDB-XXXXX.

---

**Fig. S1.**
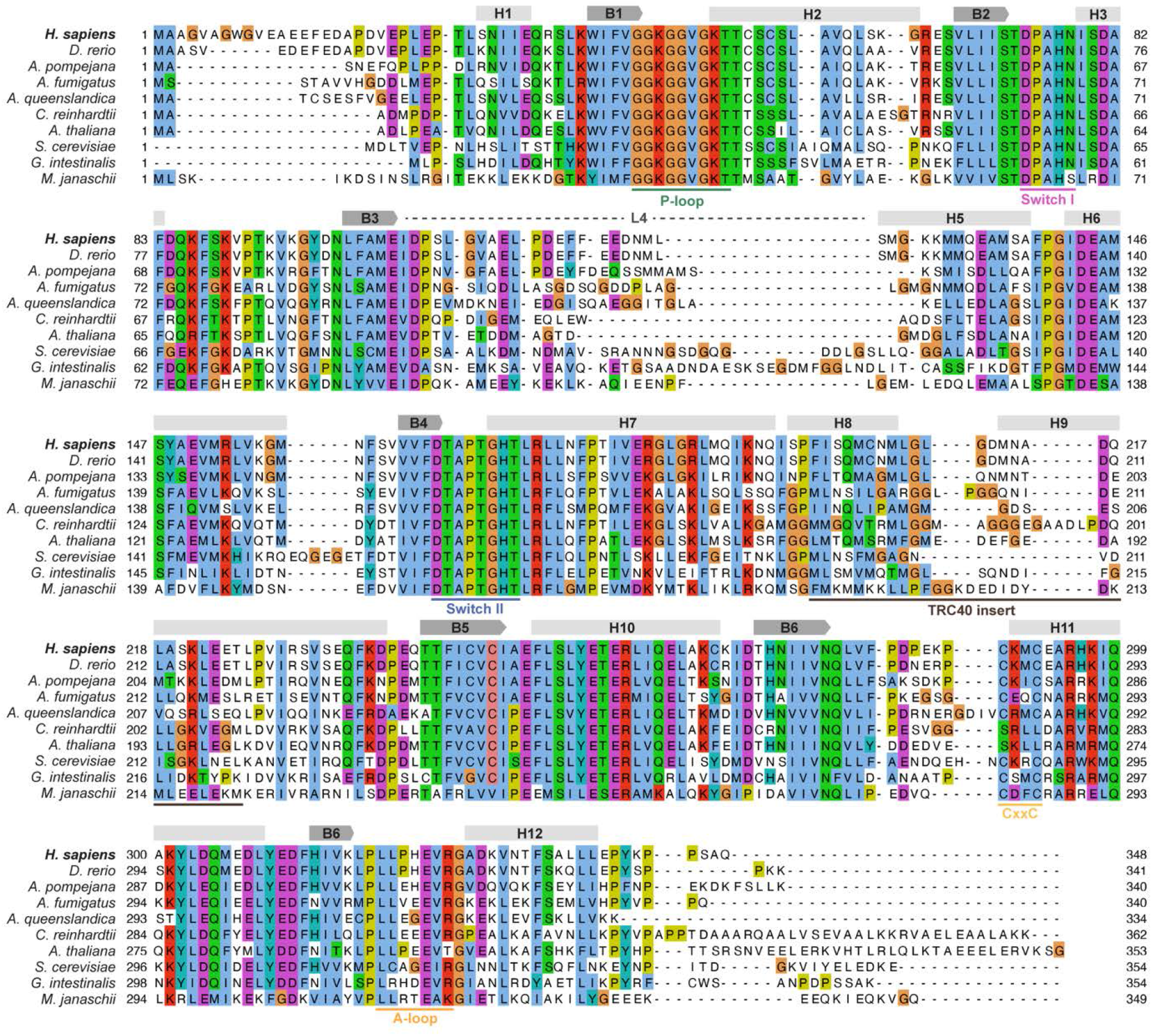
Get3 sequence alignment and annotation. Structure-based multiple sequence alignment generated using Promals3D (*71*) and visualized with Clustal coloring (*72*). Species were selected to represent phylogenetically diverse Get3 homologs. Secondary structure elements derived from the *Hs*Get3 structure are shown above the alignment, and Walker A ATPase motifs are indicated below the alignment.

**Fig. S2.**
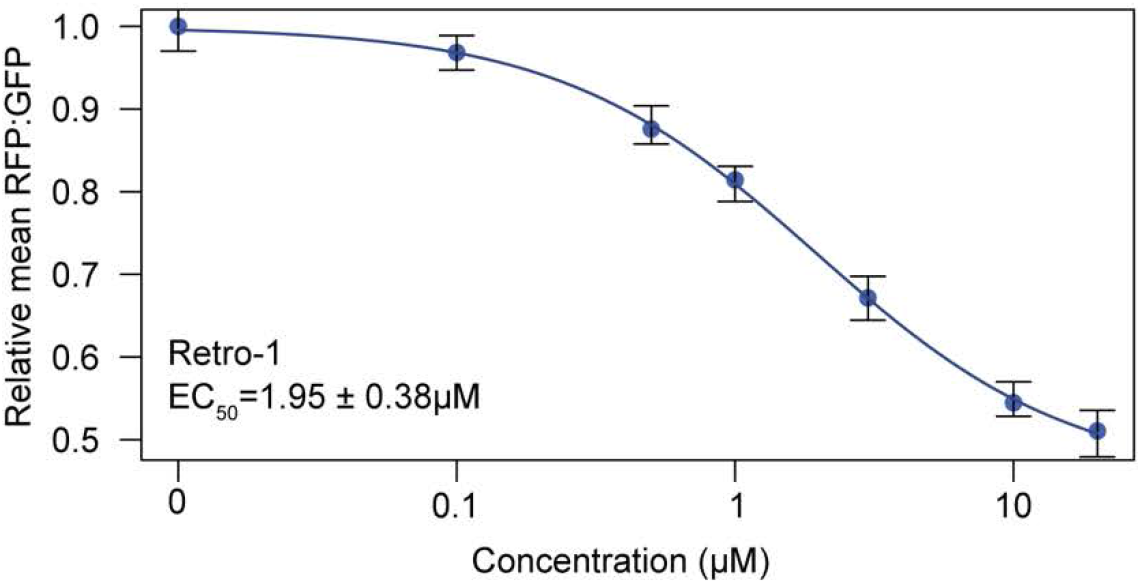
TA reporter assay dose response curve for Retro-1. Wild-type HEK293T cells expressing the GFP-2A-RFP-Sec61B_TMD_ were pre-treated with the indicated concentrations of Retro-1 for 1 h prior to induction with doxycycline for approximately 18 h, followed by FACS analysis. Shown is the dose-response curve for the reporter RFP:GFP ratios as relative means (3 experiments) to the mock-treated cells. The dose response was modeled using the four-parameter logistic regression to determine the half maximal effective concentration (EC_50_ ± standard error). Error bars for the means represent the standard error calculated from the four-parameter logistic regression.

**Fig. S3.**
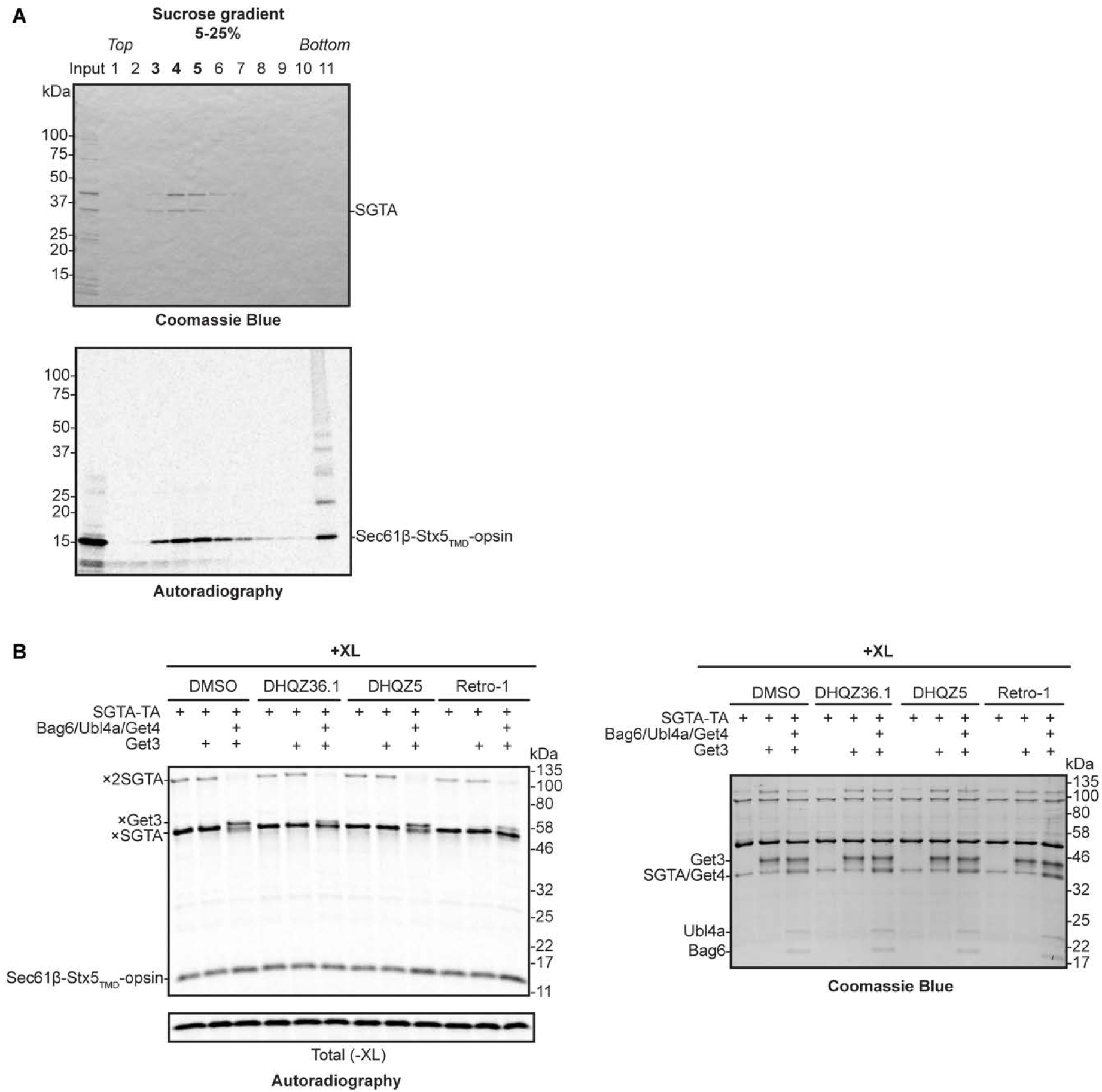
TA handover assay extended data. (**A**) SDS-PAGE analysis of 5-25% sucrose gradient separation of SGTA-TA complexes. Fractions 3-5, corresponding to SGTA-TA complexes, were pooled for use in TA handover assays. (**B**) SDS-PAGE analysis of TA handover assays after crosslinking. Autoradiography (left) and Coomassie Blue staining (right) of the same gel are shown. Figure expanded from Figure 5e of Morgens et al., 2019 (*27*)

**Fig. S4.**
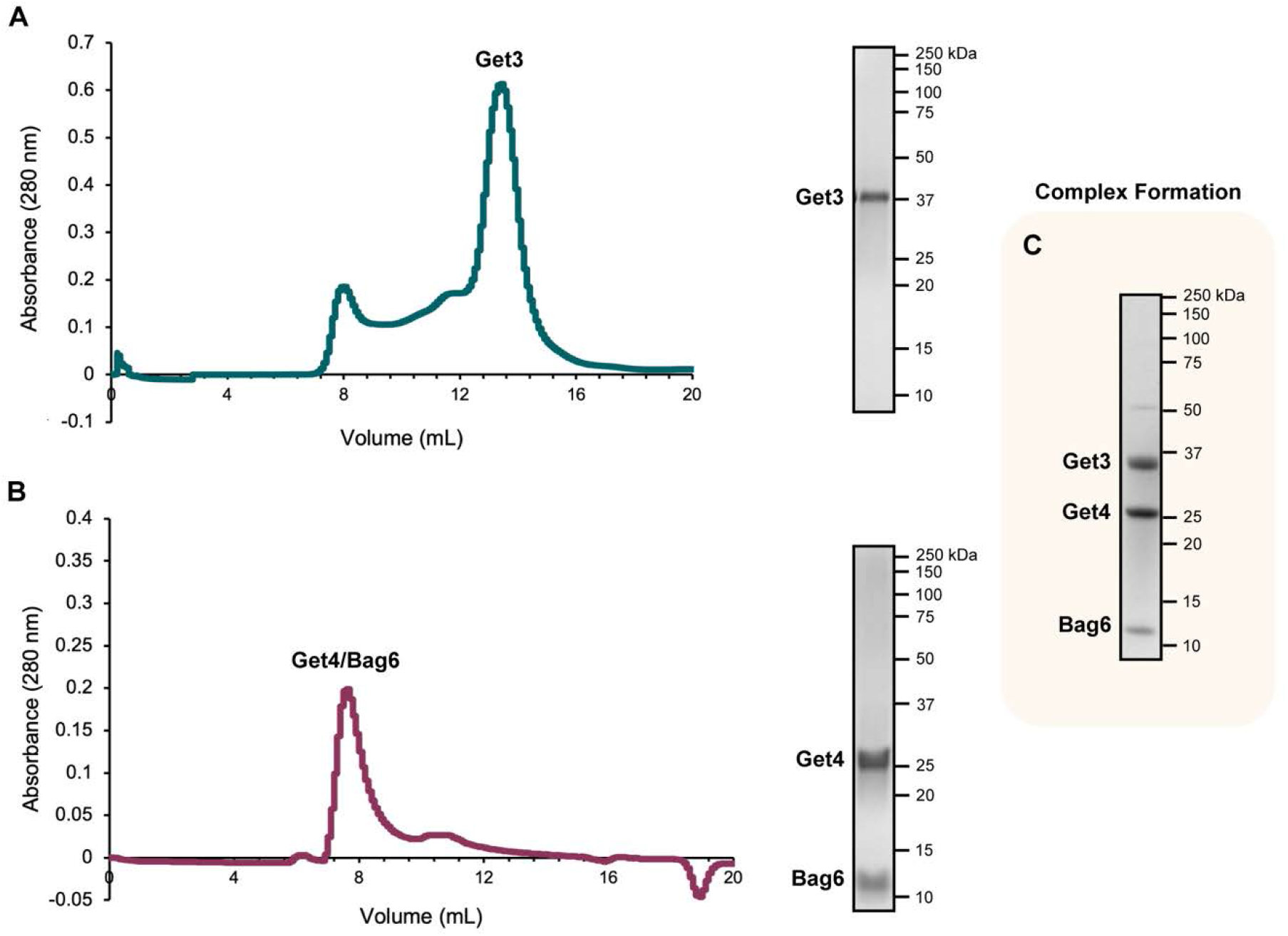
Purification of *Hs*Get3 and *Hs*Get4/Bag6. (**A**) Size-exclusion chromatogram and corresponding SDS-PAGE analysis of purified *Hs*Get3 on SS200 column. (**B**) Size-exclusion chromatogram and SDS-PAGE analysis of the copurified *Hs*Get4/Bag6 complex on SS75 column. (**C**) SDS-PAGE analysis of the *Hs*Get3–Get4/Bag6 complex used for cryo-EM preparation with Retro compounds.

**Fig. S5.**
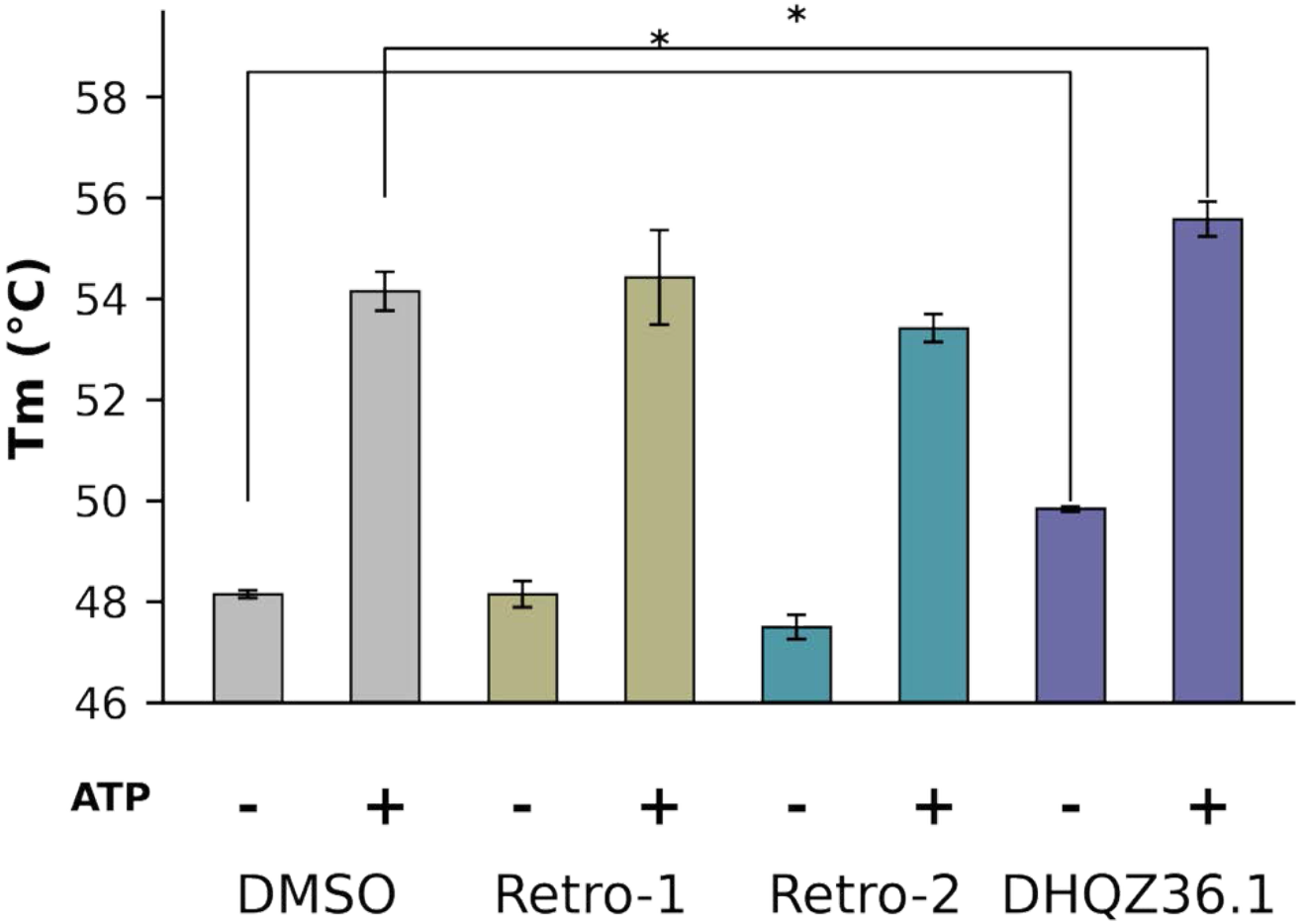
Differential scanning fluorimetry melting curves. The thermostability of the human Get3 protein was determined using differential scanning fluorimetry. Experiments were run in triplicate, with the average melting temperature indicated by the height of the bars. The standard error of the triplicates is indicated by error bars. Significance is indicated at a *p*-value of 0.05 between the DMSO control group and the experimental groups with an asterisk.

**Fig. S6.**
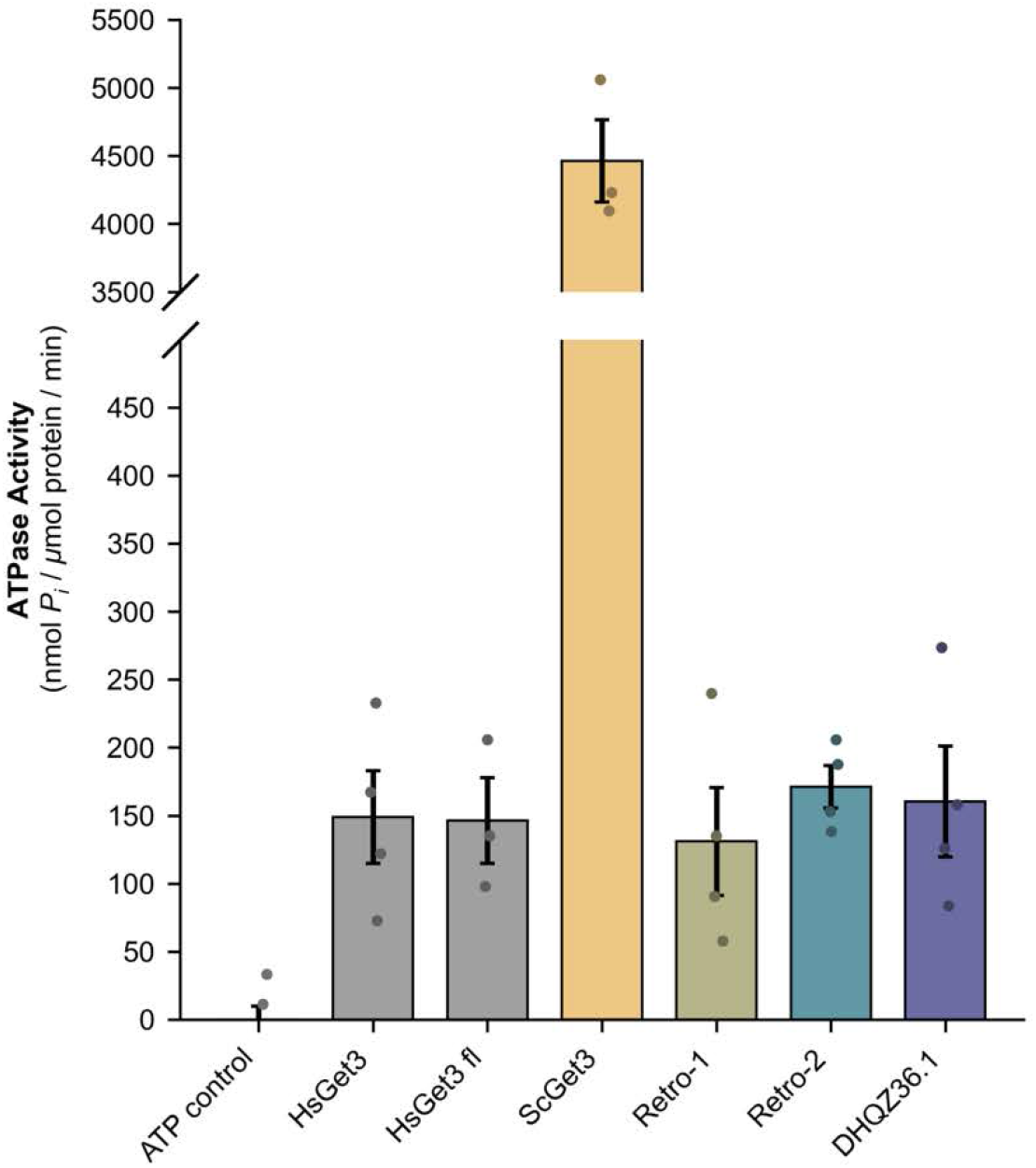
ATPase assay extended data. ATPase assays were performed using the Malachite Green Phosphate Assay Kit. Indicated compounds were added to a final concentration of 500 µM. Timepoints were measured every 15 min for 1 h, and absorbance was measured at 620 nm. Phosphate was quantified by comparison to standards. Data are shown as mean ± SE with independent replicates shown.

**Fig. S7.**
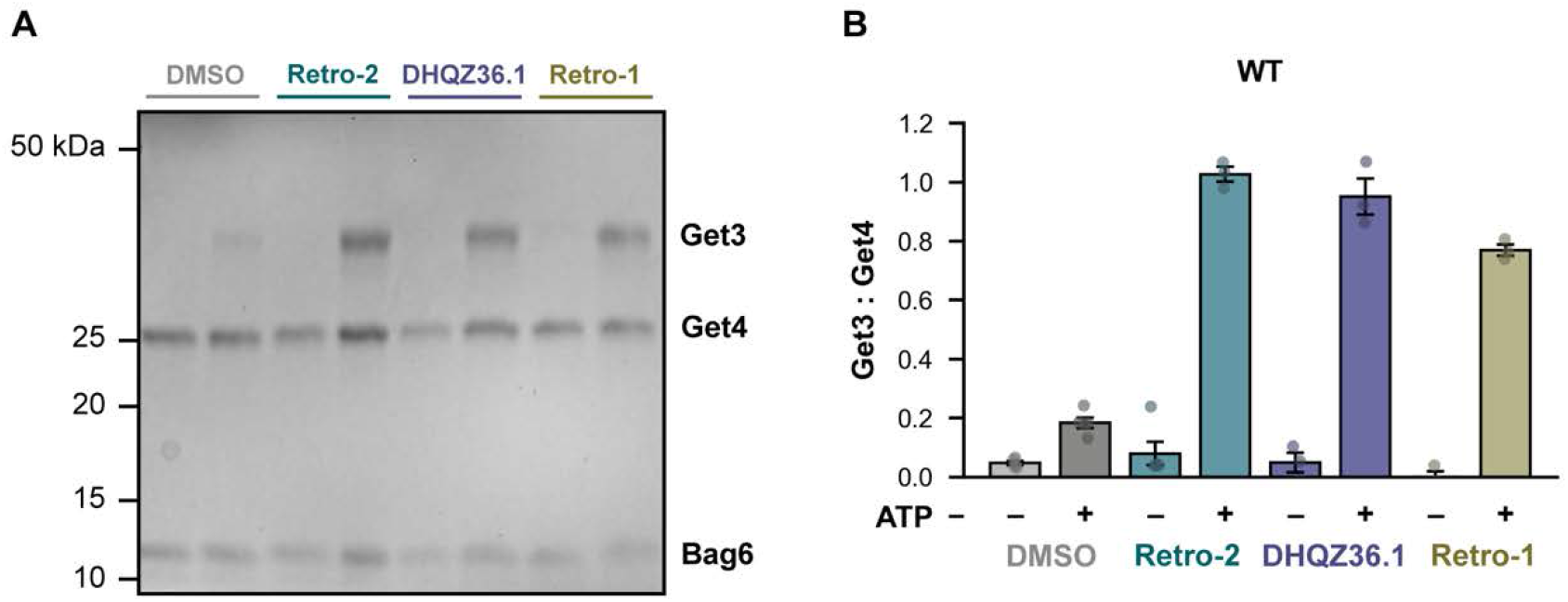
Capture assay extended data. (**A**) Coomassie-Blue-stained SDS-PAGE analysis used for quantification of Get3 pulldown by Get4/Bag6. (**B**) Quantification of capture assays performed in the presence of ATP and/or Retro compounds. Data are shown as means ± standard error with individual data points from ≥3 replicates.

**Fig. S8.**
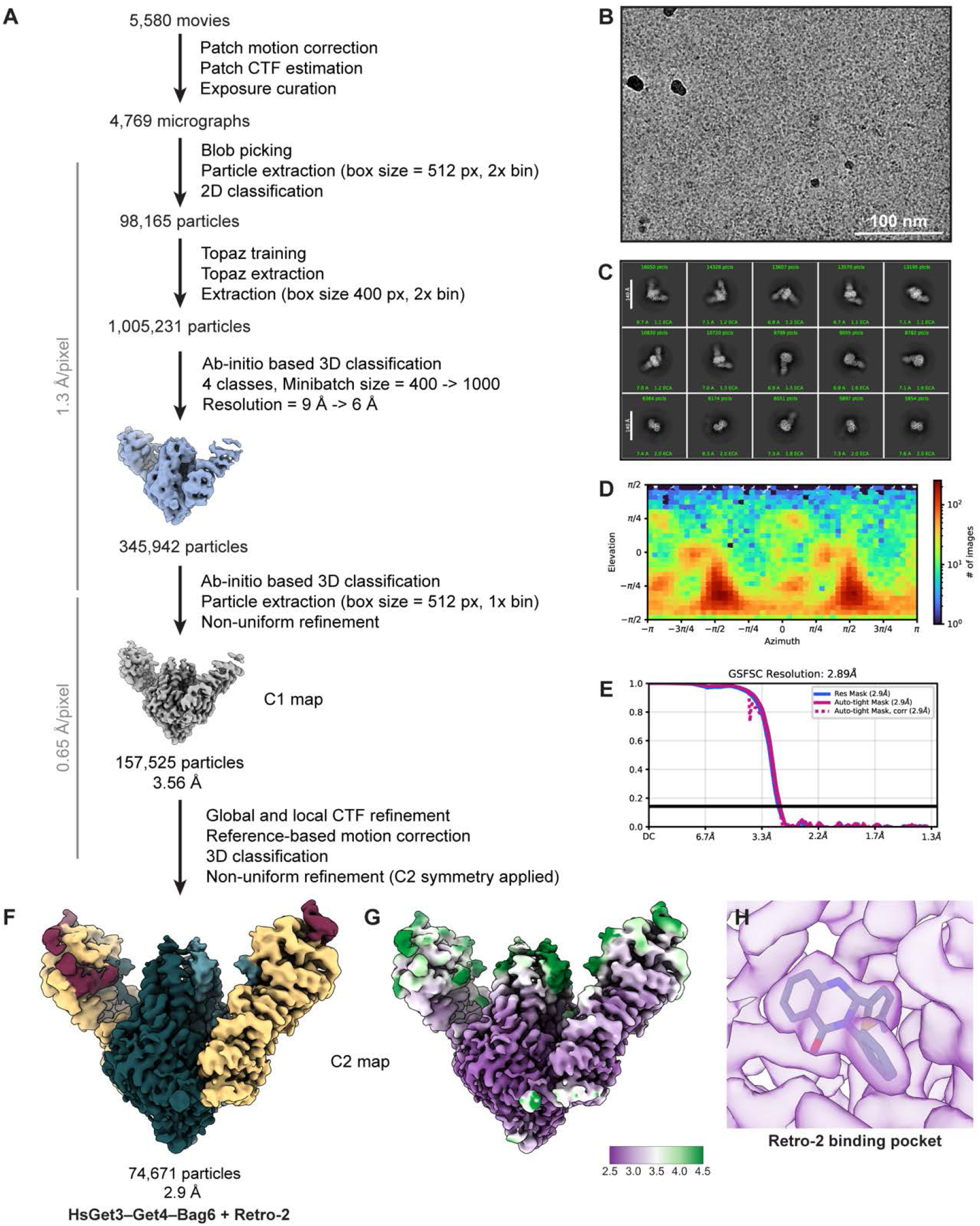
Cryo-EM data processing for Retro-2-bound *Hs*Get3–Get4/Bag6 structure. (**A**) Summary of cryo-EM data processing workflow used to obtain the Retro-2-bound complex structure. (**B**) Representative micrograph. (**C**) Representative 2D class averages. (**D**) Angular distribution of particles contributing to the final 3D reconstruction. (**E**) Fourier shell correlation (FSC) curve between the two half maps. (**F**) Cryo-EM density colored by component (Get3, teal; Get4, yellow; Bag6, magenta). (**G**) Local resolution map colored by resolution, from purple (2.5 Å) to green (4.5 Å). (**H**) Local resolution at Retro-2 binding pocket, colored as in (G).

**Fig. S9.**
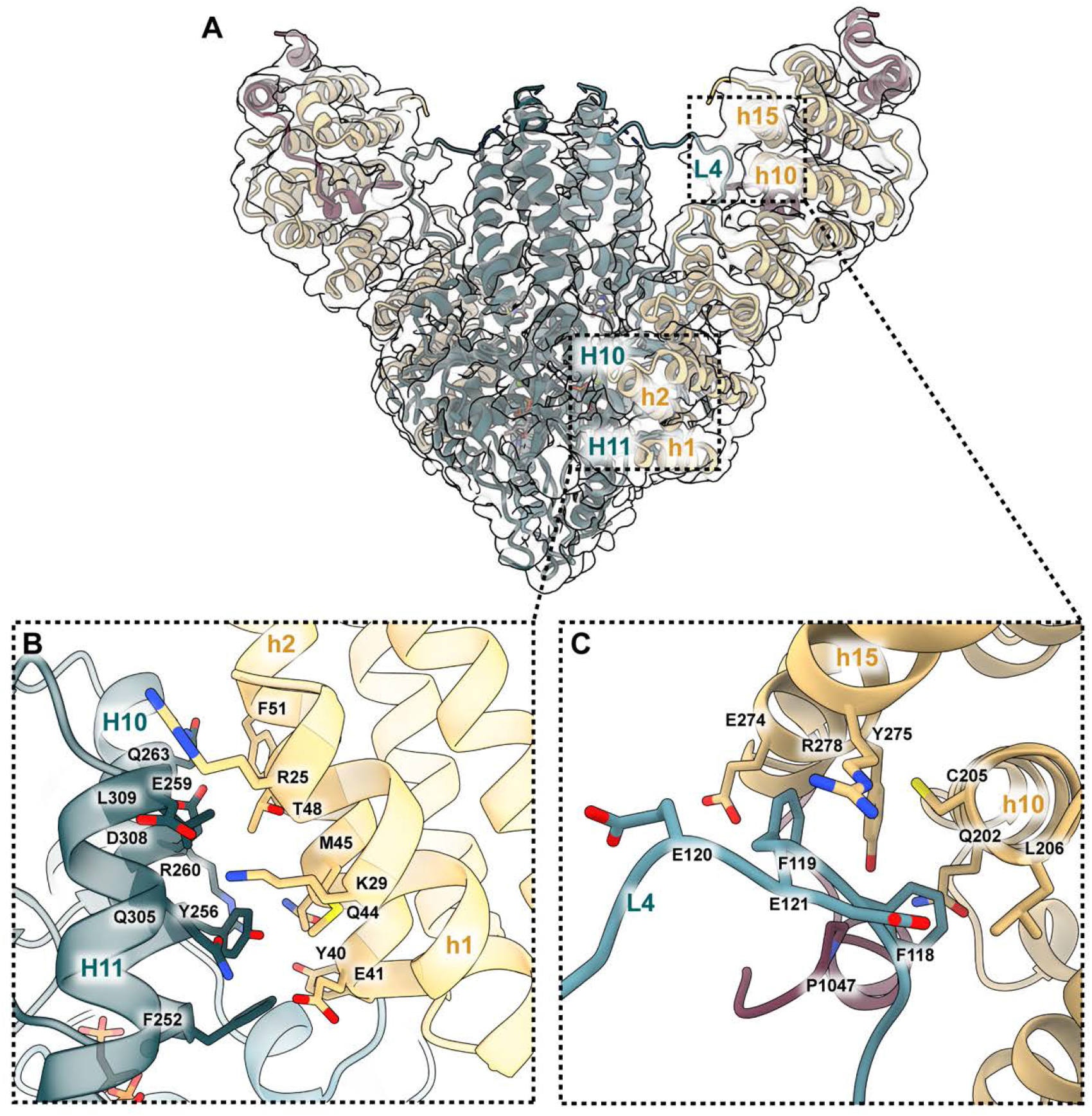
Get3–Get4/Bag6 interaction interfaces. (**A**) Retro-2 bound Get3–Get4/Bag6 model shown inside of cryo-EM density, with interacting interfaces labeled. (**B**) Close-up of Get3 H10/H11 interaction with Get4 h2/h1, with pertinent residues shown in sticks and labeled. (**C**) Close-up of Get3 L4 interaction with Get4 h10/h15 and C-terminus of Bag6, with pertinent residues show in sticks and labeled.

**Fig. S10.**
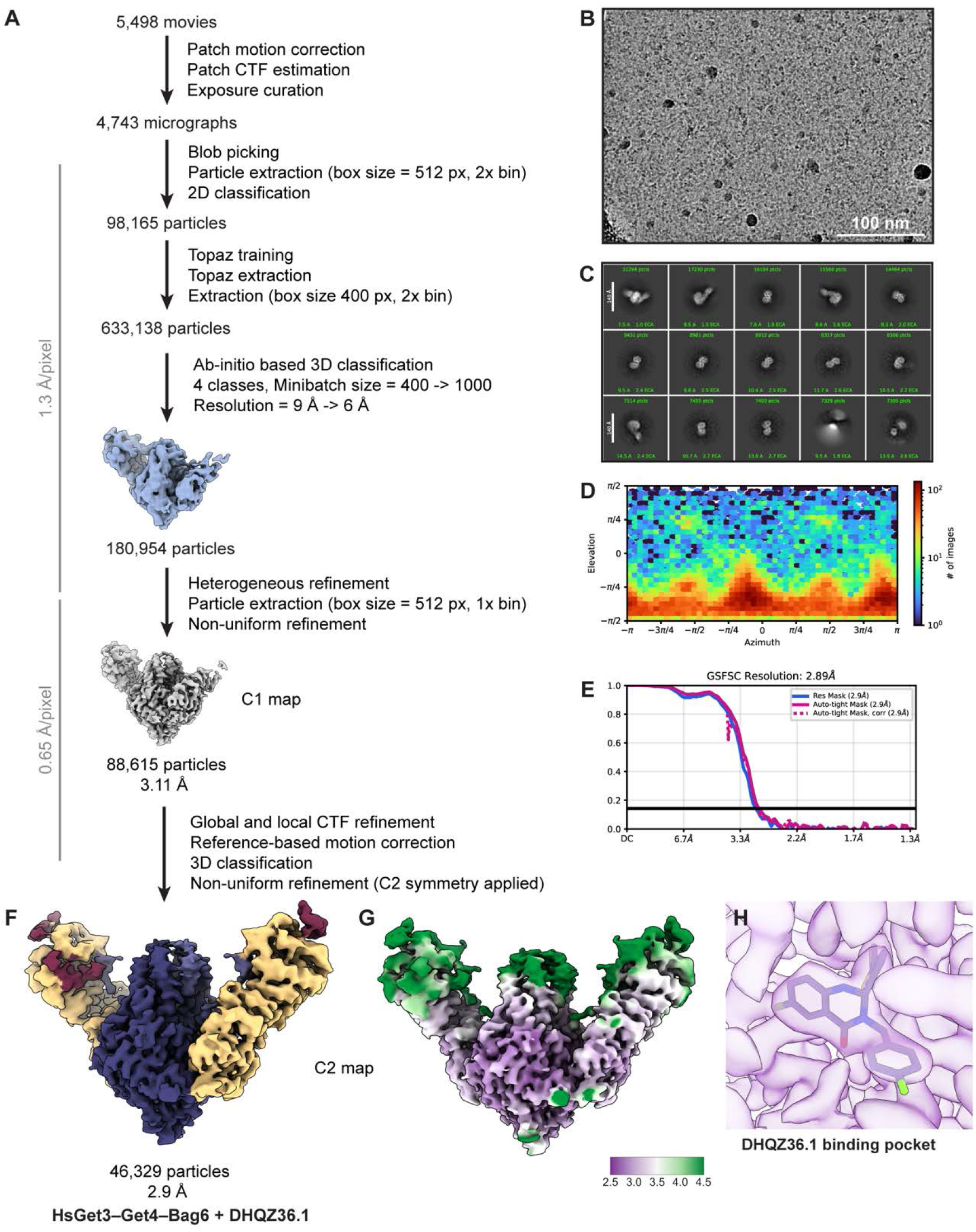
Cryo-EM data processing for DHQZ36.1-bound *Hs*Get3–Get4/Bag6 structure. (**A**) Summary of cryo-EM data processing workflow used to obtain the DHQZ36.1-bound complex structure. (**B**) Representative micrograph. (**C**) Representative 2D class averages. (**D**) Angular distribution of particles contributing to the final 3D reconstruction. (**E**) Fourier shell correlation (FSC) curve between the two half maps. (**F**) Cryo-EM density colored by component (Get3, purple; Get4, yellow; Bag6, magenta). (**G**) Local resolution map colored by resolution, from purple (2.5 Å) to green (4.5 Å). (**H**) Local resolution at DHQZ36.1 binding pocket, colored as in (G).

**Fig. S11.**
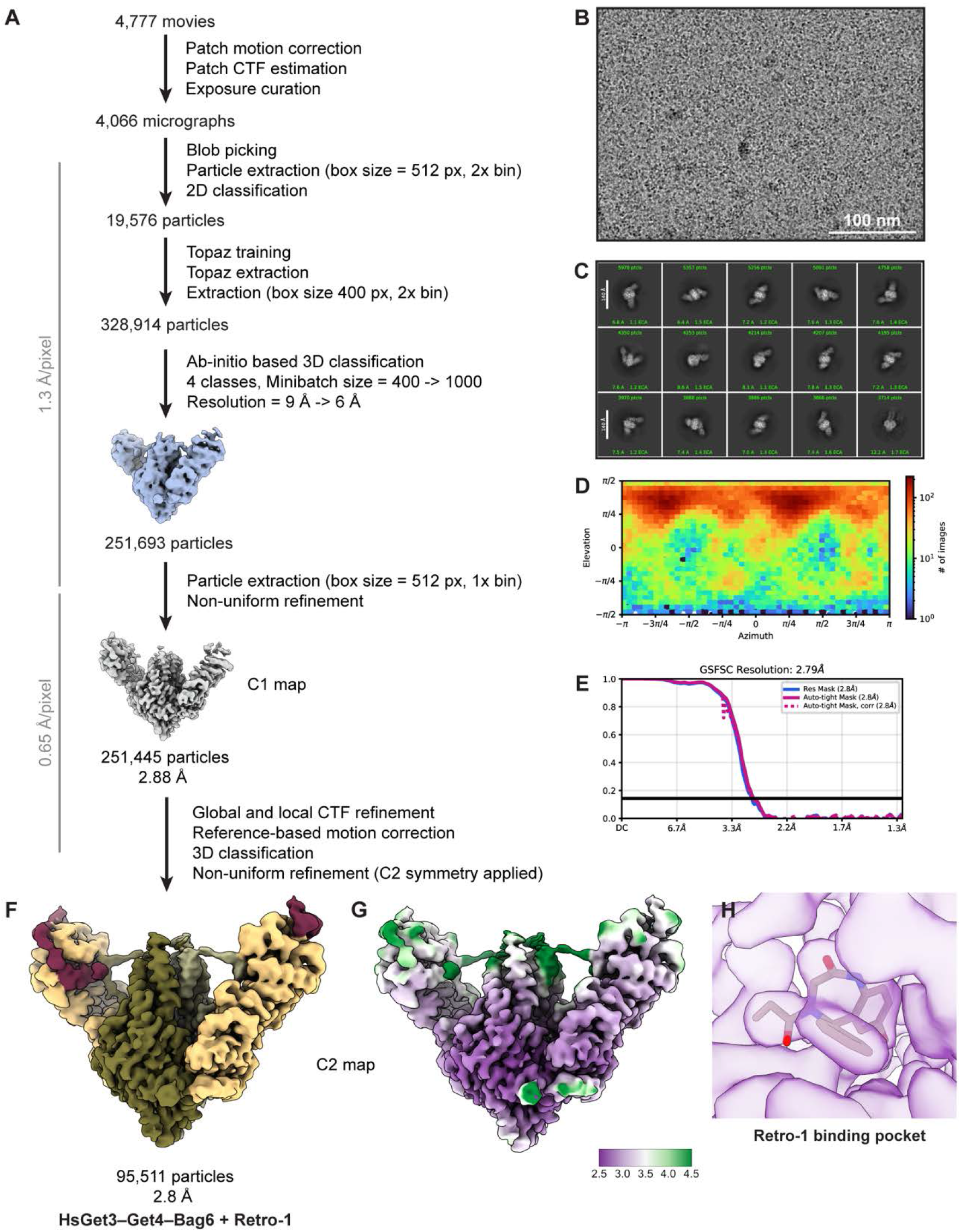
Cryo-EM data processing for Retro-1-bound *Hs*Get3–Get4/Bag6 structure. (**A**) Summary of cryo-EM data processing workflow used to obtain the Retro-1-bound complex structure. (**B**) Representative micrograph. (**C**) Representative 2D class averages. (**D**) Angular distribution of particles contributing to the final 3D reconstruction. (**E**) Fourier shell correlation (FSC) curve between the two half maps. (**F**) Cryo-EM density colored by component (Get3, green; Get4, yellow; Bag6, magenta). (**G**) Local resolution map colored by resolution, from purple (2.5 Å) to green (4.5 Å). (**H**) Local resolution at Retro-1 binding pocket, colored as in (G).

**Fig. S12.**
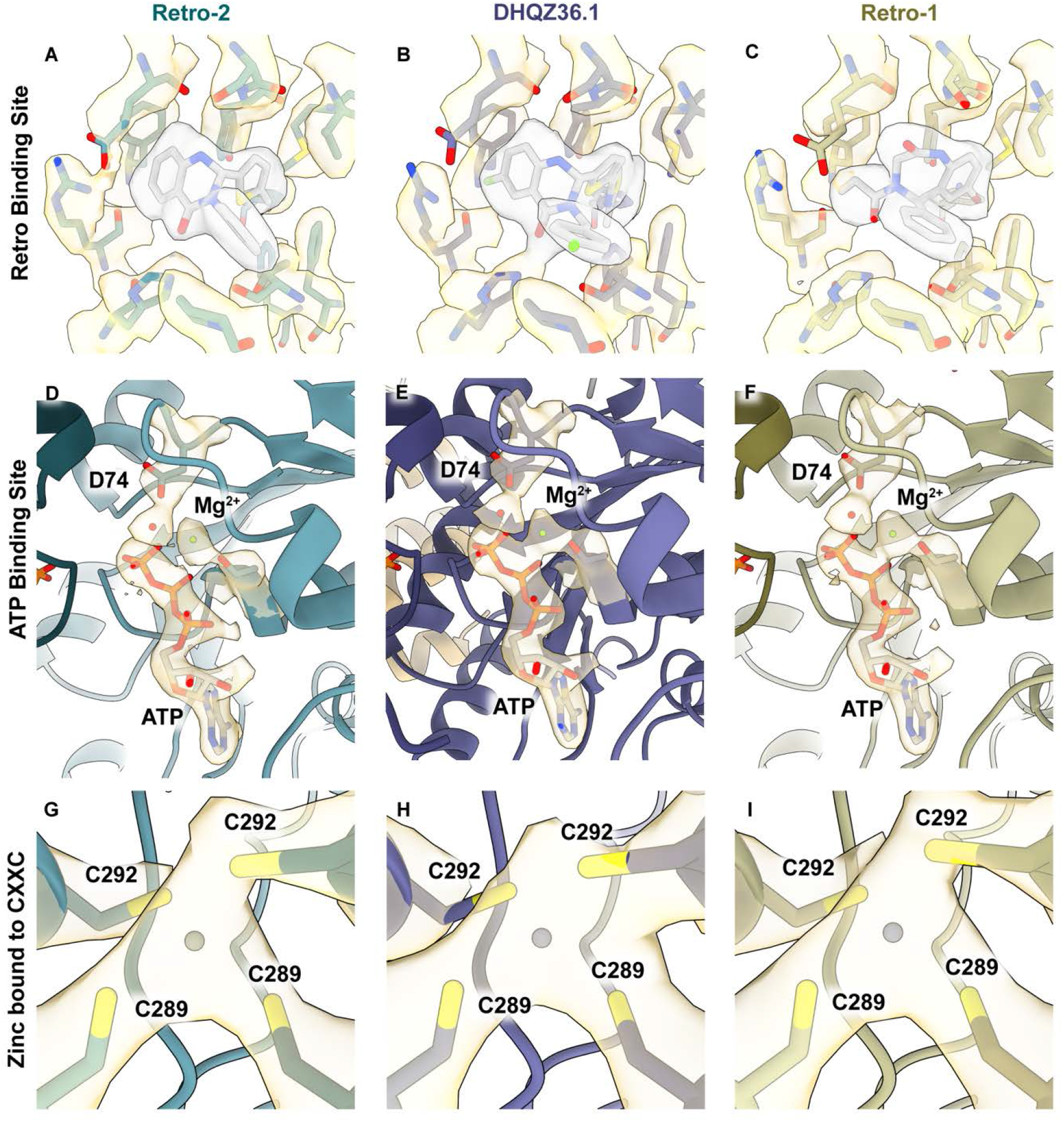
Density for key motifs in Retro-bound *Hs*Get3–Get4/Bag6 complex. (**A**) Cryo-EM density showing fitted Retro-2 (grey) and binding residues (yellow). (**B**) Cryo-EM density showing fitted DHQZ36.1 (grey) and binding residues (yellow). (**C**) Cryo-EM density showing fitted Retro-1 (grey) and binding residues (yellow). (**D**) Cryo-EM density showing fitted ATP, Mg^2+^, coordinating water molecule, and catalytic Asp74 in Retro-2-bound complex. (**E**) Cryo-EM density showing fitted ATP, Mg^2+^, coordinating water molecule, and catalytic Asp74 in DHQZ36.1-bound complex. (**F**) Cryo-EM density showing fitted ATP, Mg^2+^, coordinating water molecule, and catalytic Asp74 in Retro-1-bound complex. (**G**) Cryo-EM density showing fitted Zn^2+^ coordinated by four cysteines, two from each monomer (Cys289 and Cys292), for Retro-2-bound complex. (**H**) Cryo-EM density showing fitted Zn^2+^ coordinated by four cysteines for DHQZ36.1-bound complex. (**I**) Cryo-EM density showing fitted Zn^2+^ coordinated by four cysteines for Retro-1-bound complex.

**Fig. S13.**
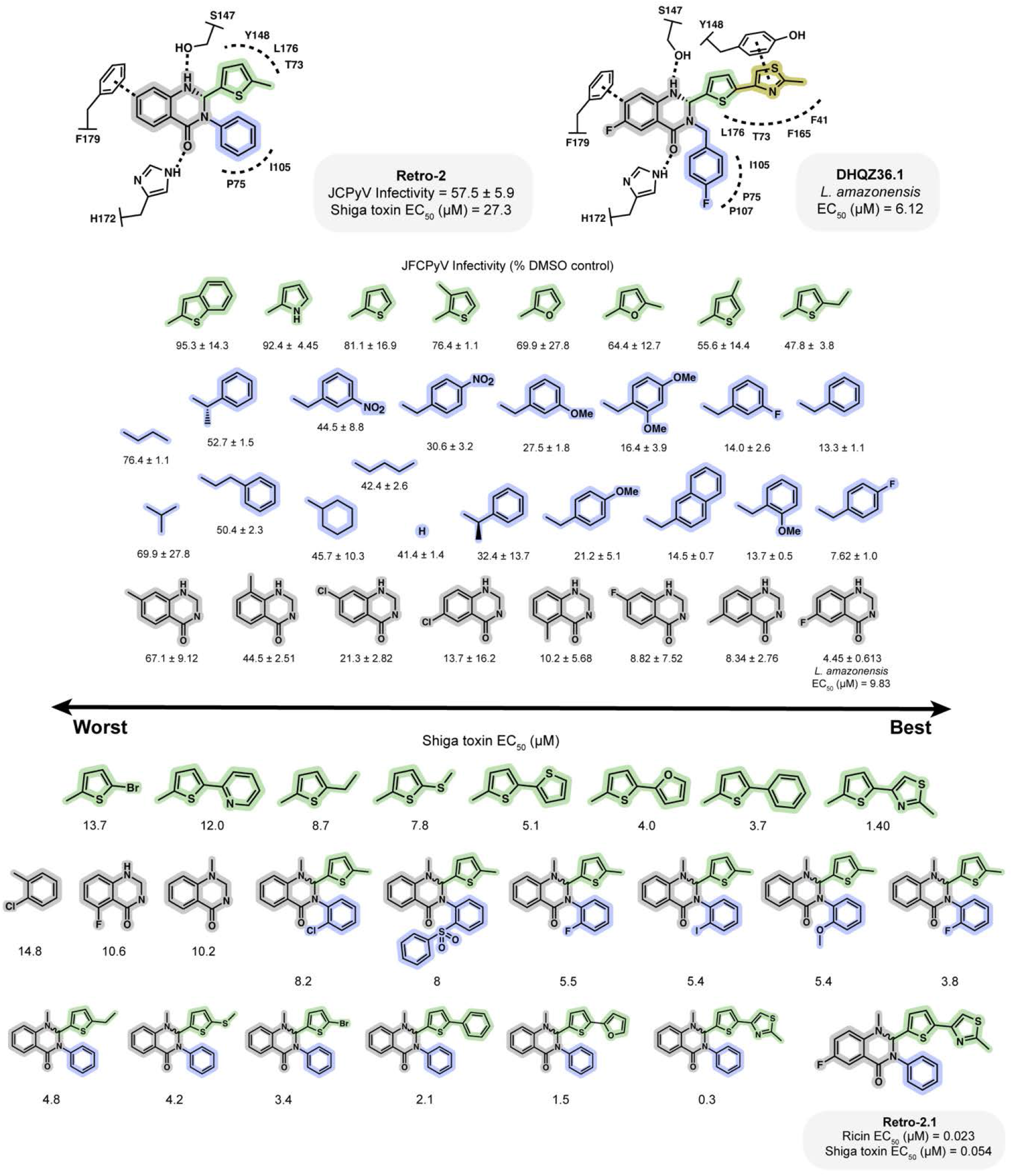
Structure–activity relationship (SAR) analysis of Retro-2. Summary of Retro-2 SAR data compiled from Gupta et al. 2013 (*13*), Noel et al. 2013 (*52*), Carney et al. 2014 (*15*), Craig et al. 2017 (*14*). and Substituent modifications are indicated, along with infectivity against JC polyomavirus after normalization to a DMSO-treated control, and/or EC_50_ values (µM) measured against *Leishmania amazonensis* or Shiga toxin/ricin toxin.

**Fig. S14.**
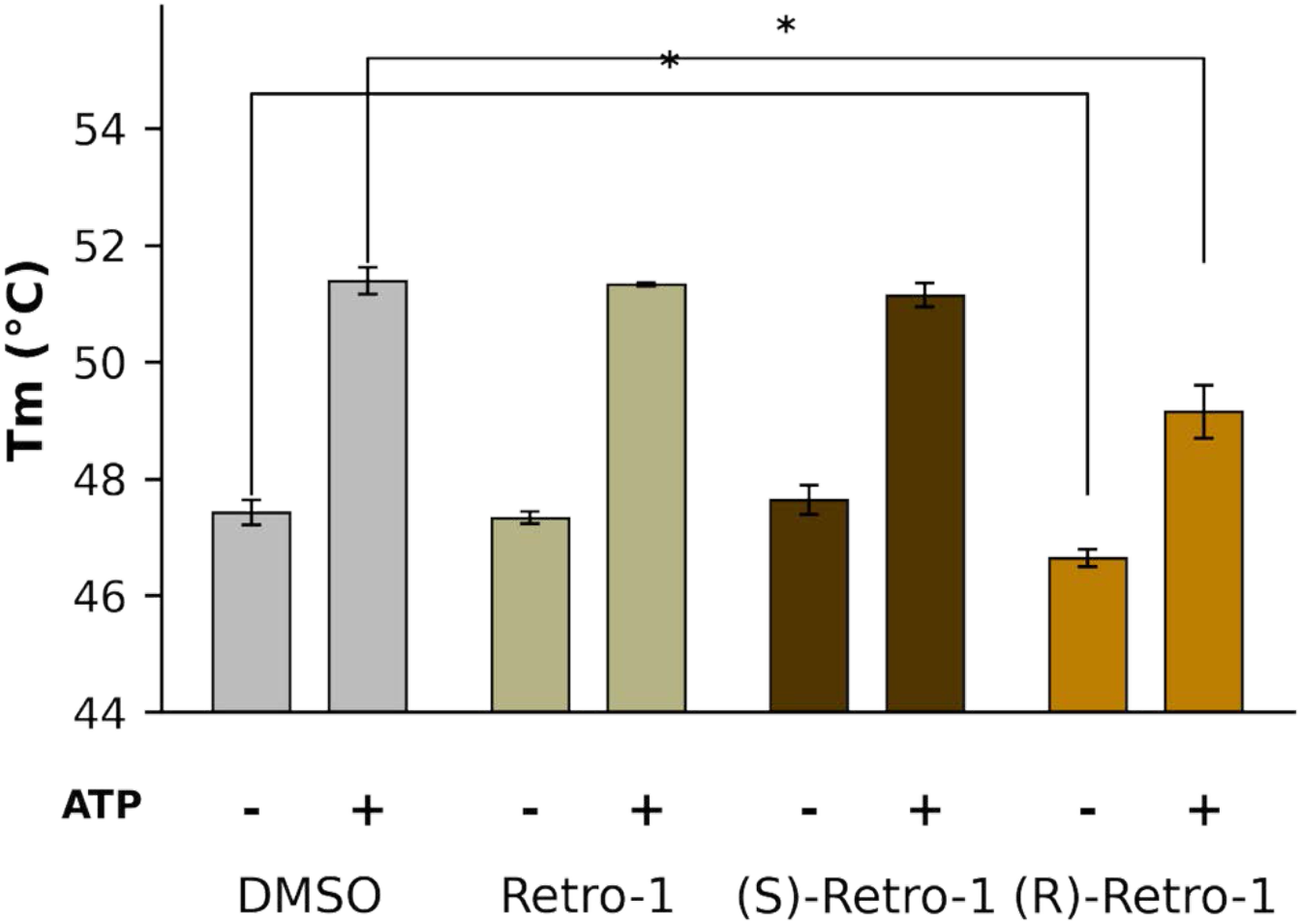
Retro-1 enantiomer differential scanning fluorimetry. Differential scanning fluorimetry was used to test the effect of the racemic mixture of Retro-1 as well as its separated enantiomers on the thermostability of human Get3. Each condition was run in triplicate, and the average melting temperature is represented by the height of the bar. The error bars represent the standard error of the triplicate. Significance is indicated at a *p*-value of 0.05 with an asterisk.

**Fig. S15.**
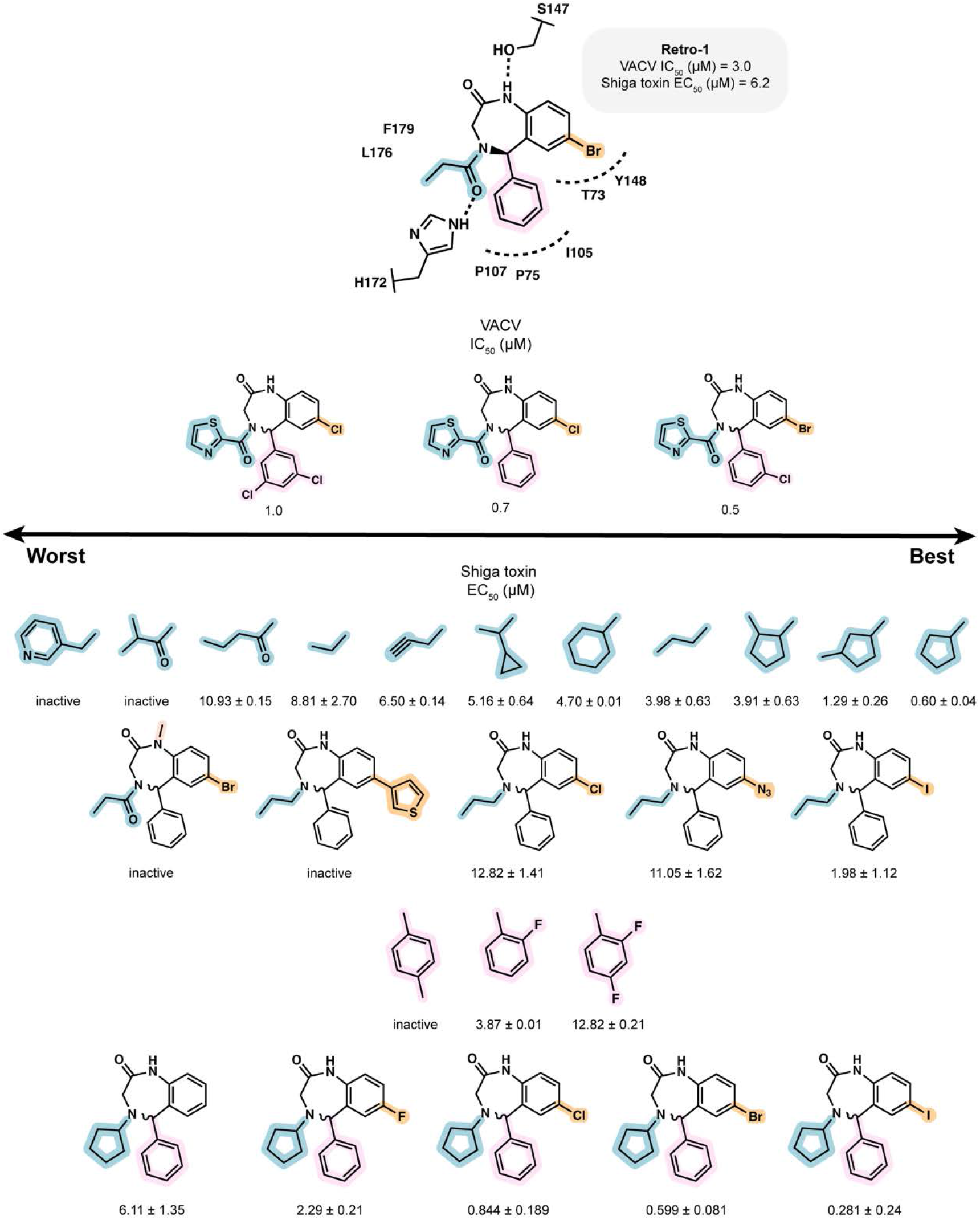
Structure–activity relationship (SAR) analysis of Retro-1. Summary of Retro-1 SAR data compiled from Abdelkafi 2020 (*11*) and Priyamvada et al. 2020 (*22*). Substituent modifications are indicated, along with IC_50_ values (µM) measured for inhibition of vaccinia virus or EC_50_ values (µM) measured against Shiga toxin.

**Fig. S16.**
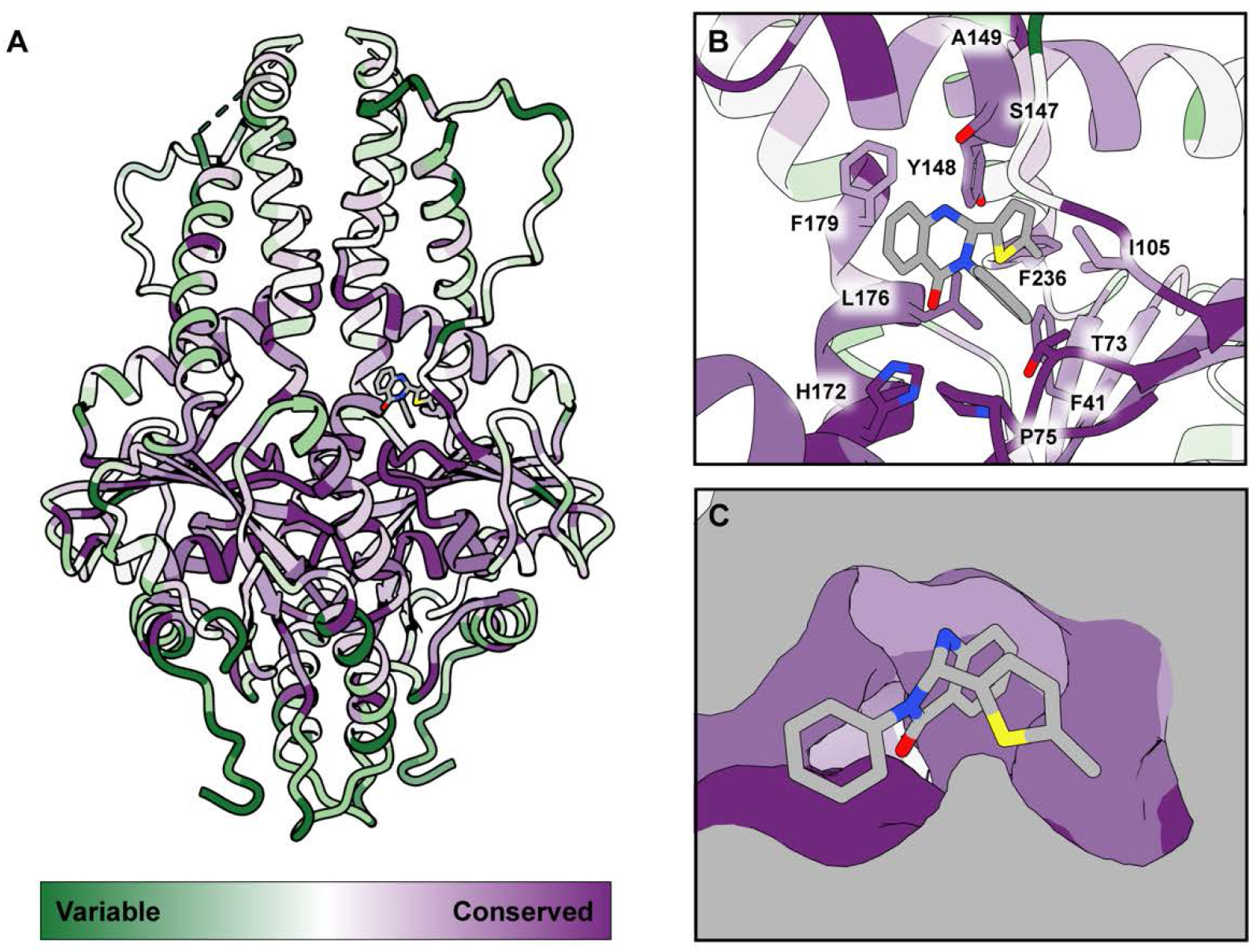
Conservation of Retro compound binding pocket. (**A**) *Hs*Get3 model from the Retro-2-bound complex shown in licorice, colored by sequence conservation (green = variable, purple = conserved) using ConSurf (*73*). (**B**) Close-up of the Retro-2 binding site from (A), with binding residues depicted as sticks and colored by conservation as in (A). (**C**) Surface representation of the Retro-2 binding, colored by conservation as in (A).

**Fig. S17.**
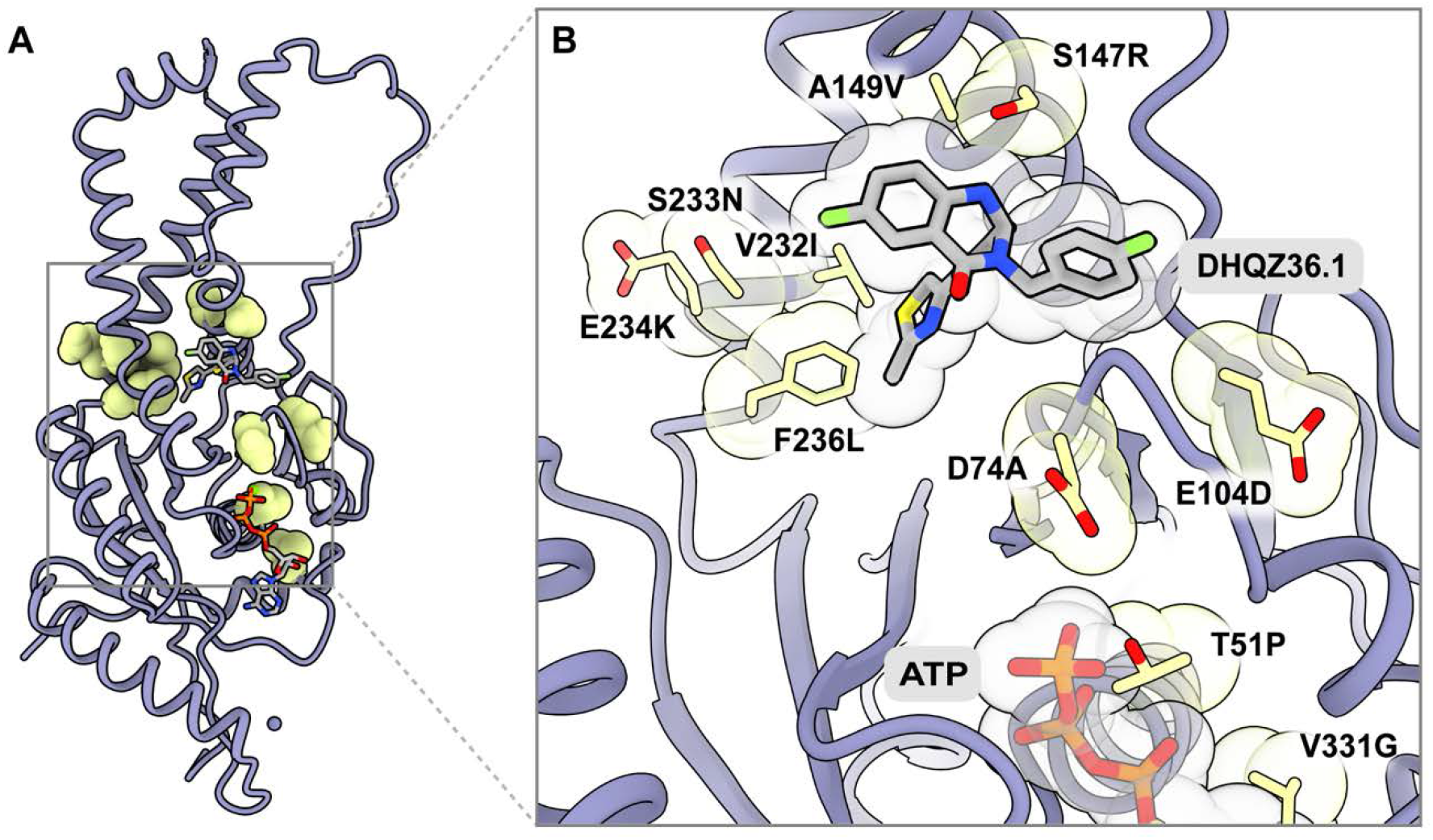
Resistant Get3 mutations found in CRISPR-X screen with DHQZ36.1. (**A**) CRISPR-X screen mutations found in Morgens et al. (*23*) to be resistant to DHQZ36.1 are shown in sphere representation, colored lime. (**B**) Close-up of the DHQZ36.1 binding site from (A), with CRISPR-X screen mutations highlighted. Helix 7 has been omitted for clarity.

**Fig. S18.**
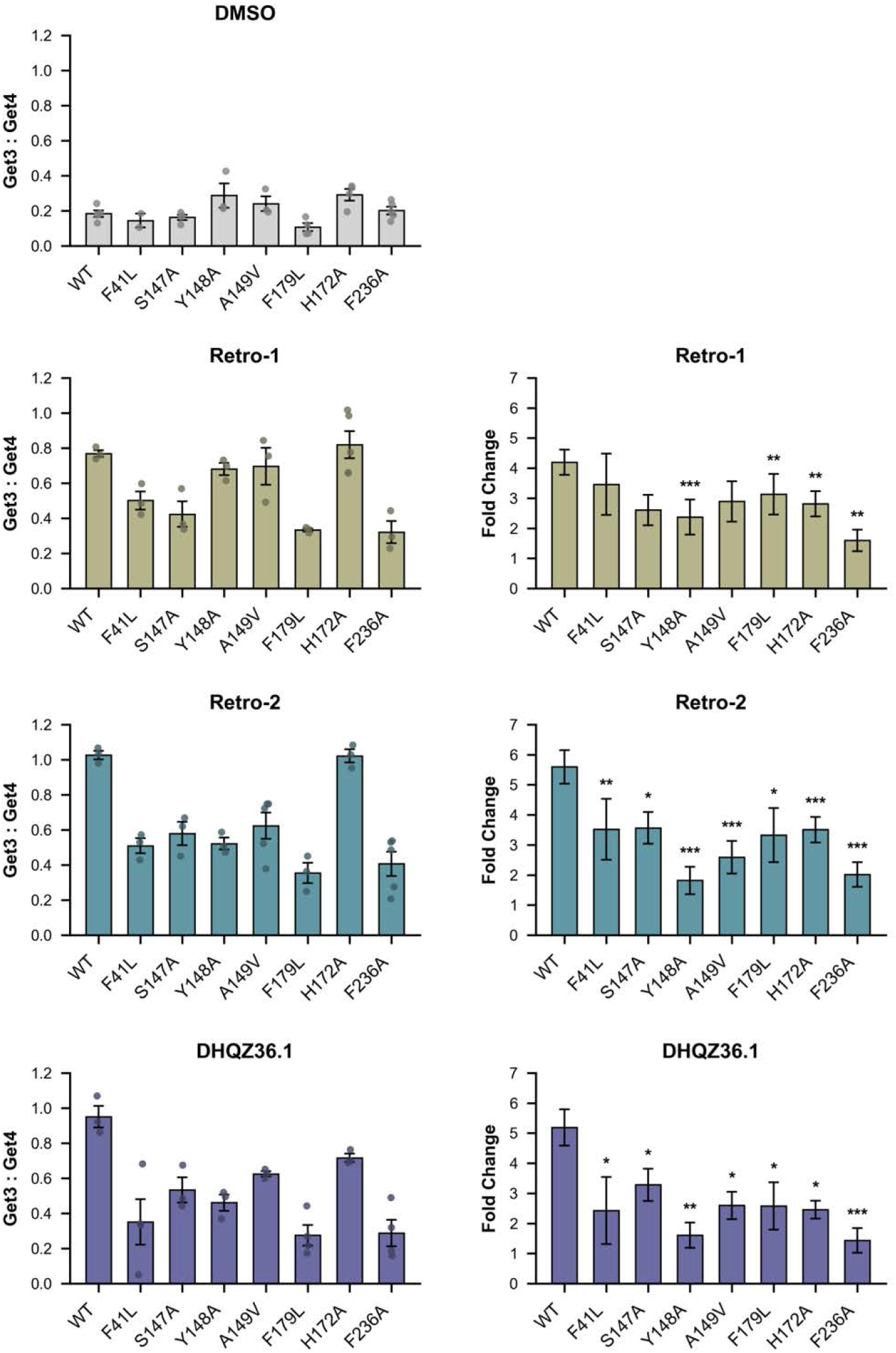
*Hs*Get3 point mutant capture assay extended data. Capture assays were performed with *Hs*Get3 point mutants in the presence of ATP. Raw data are shown as mean ± SE with individual data points from ≥ 3 replicates also shown for DMSO control, Retro-1, Retro-2, and DHQZ36.1. Fold changes were calculated for each compound-treated condition relative to WT in the presence of ATP using Welch’s t-test (statistical significance: * = *p* < 0.05*, \*\** = *p < 0.01, *** = p* < 0.001).

**Fig. S19.**
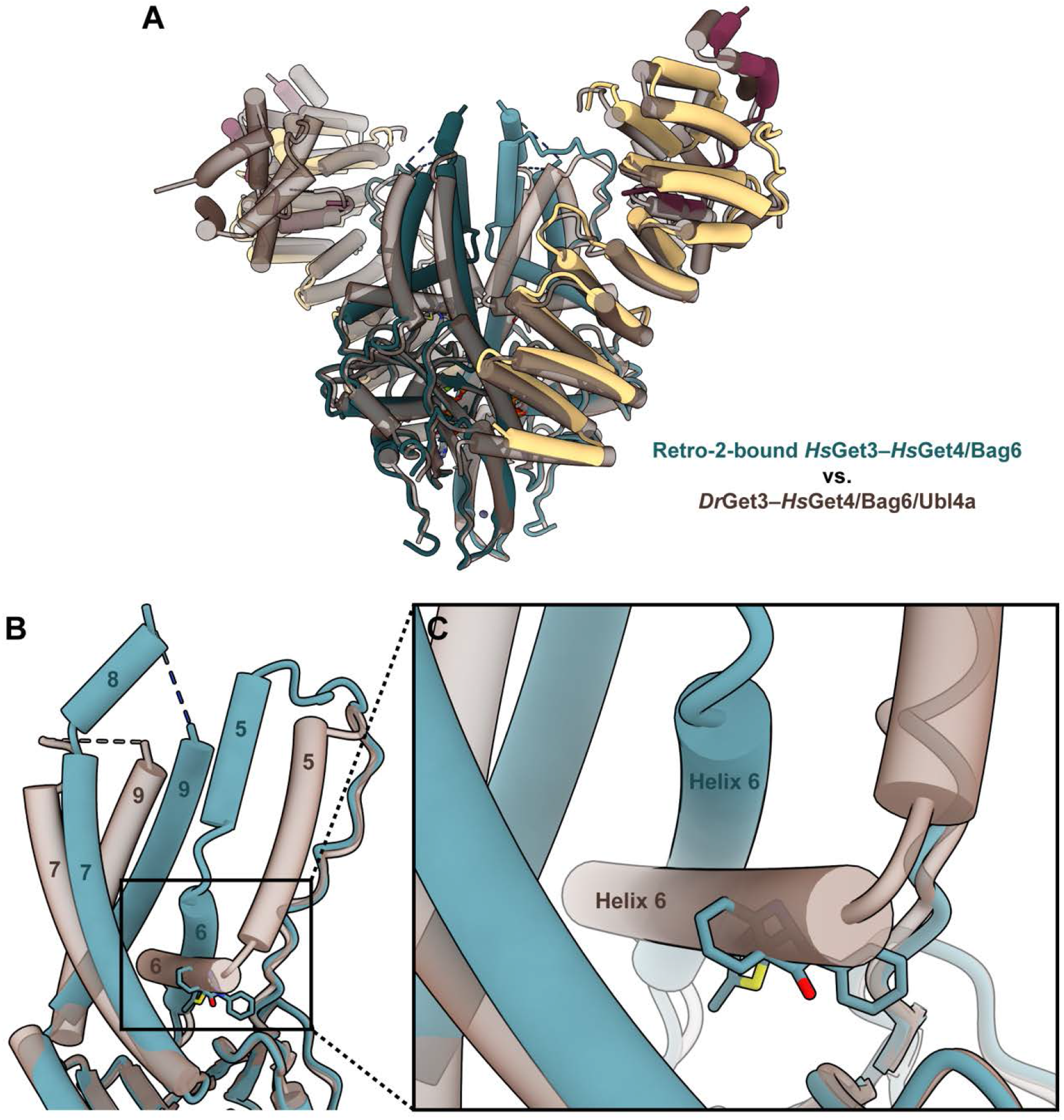
Retro-2 impedes canonical helix 6 movement in the pretargeting complex. (**A**) Comparison of Retro-2-bound *Hs*Get3–*Hs*Get4/Bag6 complex (teal) to the chimeric metazoan pretargeting *Dr*Get3–*Hs*Get4/Bag6/Ubl4a complex (brown, PDB ID: 7ru9). (**B**) Close-up of the comparison showing a single Get3 monomer with client-binding helices labeled. (**C**) Close-up of the comparison showing the positioning of Get3 H6 relative to the Retro-2 binding site.

**Fig. S20.**
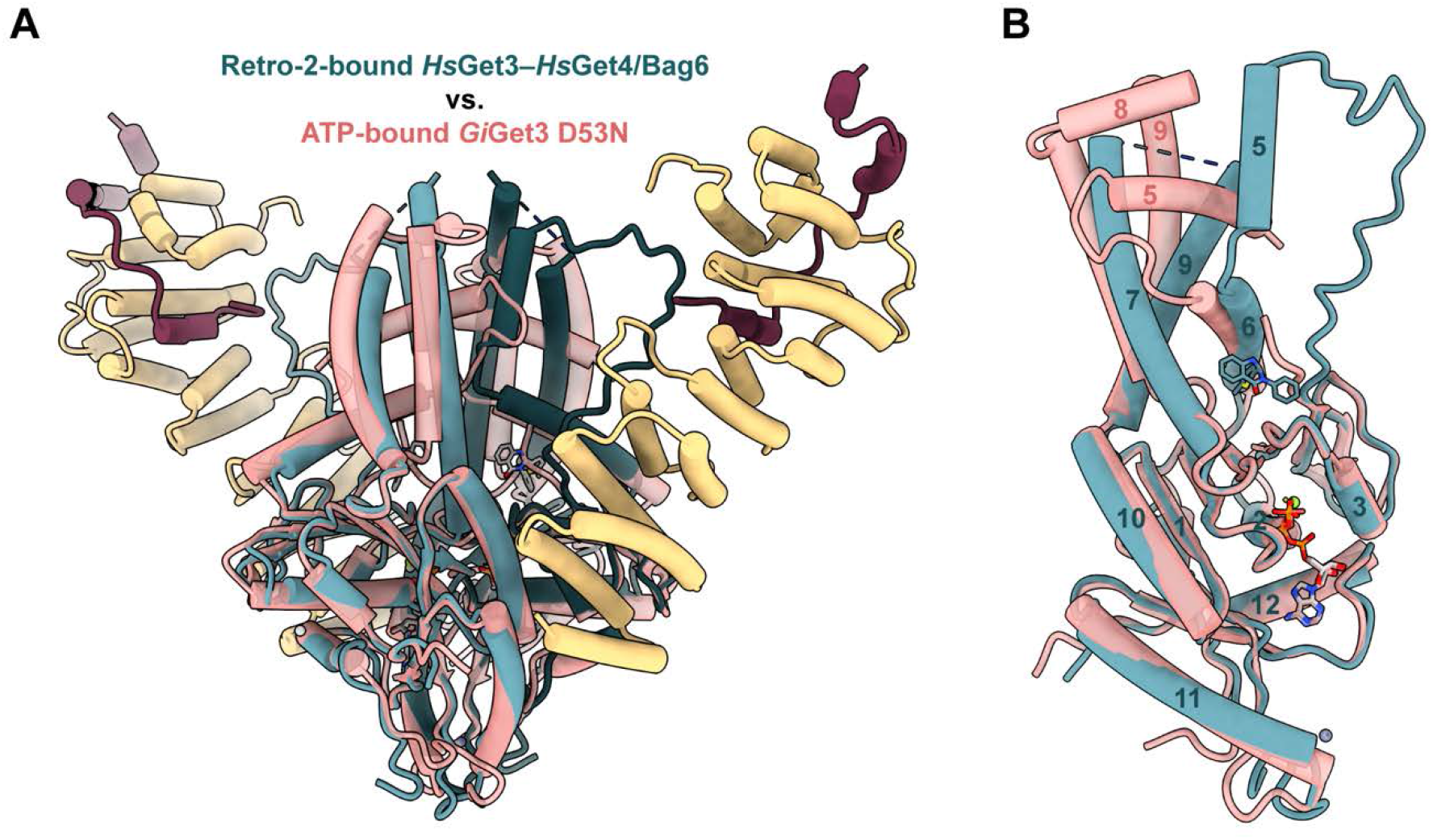
Get3 helix 6 remains in upward position despite Get4/Bag6 binding. (**A**) Comparison of Retro-2-bound *Hs*Get3–Get4/Bag6 complex (teal) to ATP-bound *Gi*Get3 D53N (transparent peach, PDB ID: 7spy). (**B**) Close-up of the comparison showing a single Get3 monomer with helices labeled.

**Table S1.**
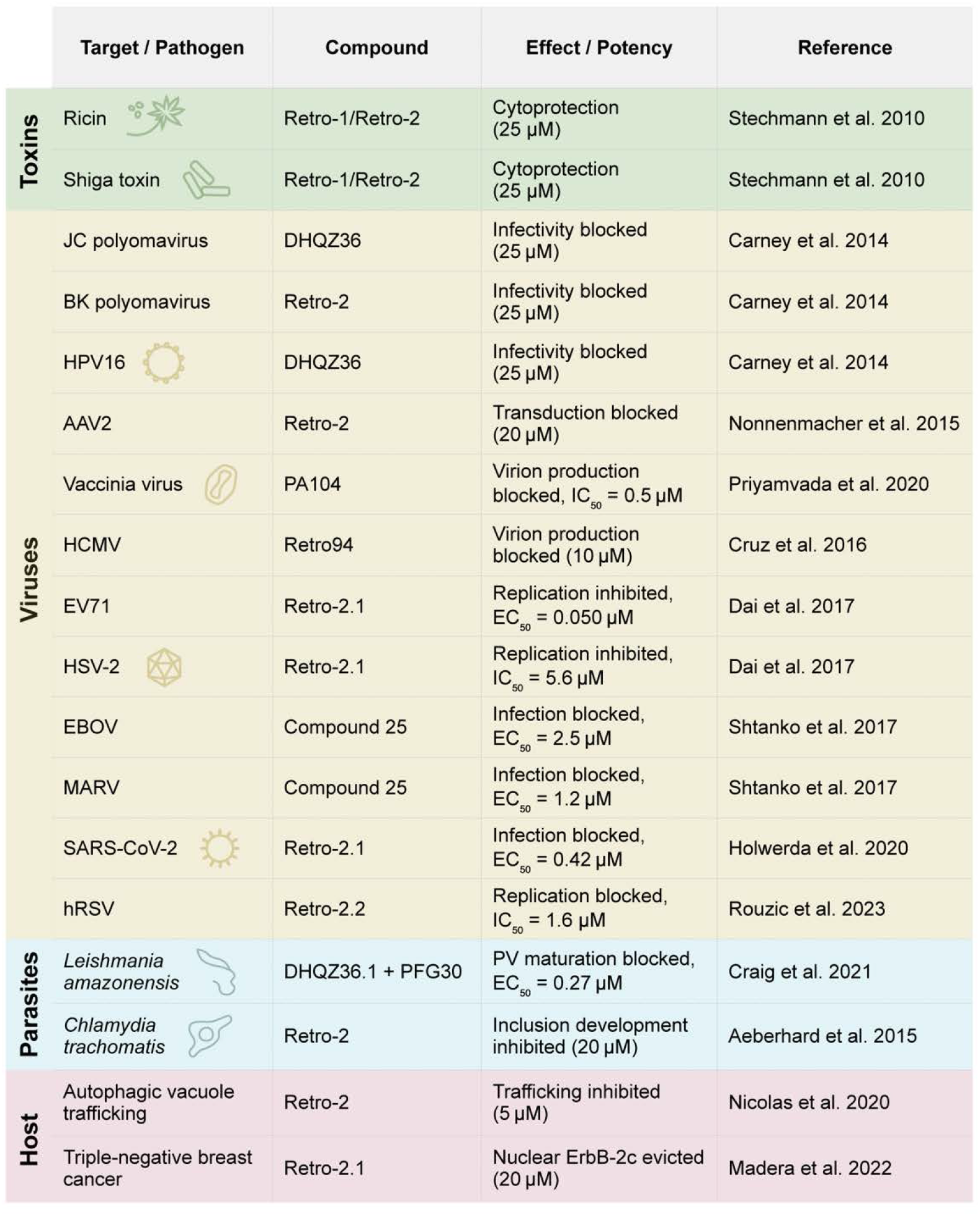
Summary of reported effects of Retro compounds.

**Table S2.** Cryo-EM data collection, refinement, and validation statistics.

|  | ***HsGet3–Get4/Bag6***<br>***+ Retro-2***<br>(EMDB-77293)<br>(PDB 35YZ) | ***HsGet3–Get4/Bag6***<br>***+ DHQZ36.1***<br>(EMDB-77294)<br>(PDB 35ZA) | ***HsGet3–Get4/Bag6***<br>***+ Retro-1***<br>(EMDB-77295)<br>(PDB 35ZB) |
| --- | --- | --- | --- |
| ***Data collection and processing*** |  |  |  |
| Microscope | FEI Titan Krios | FEI Titan Krios | FEI Titan Krios |
| Magnification | 130,000 | 130,000 | 130,000 |
| Voltage (kV) | 300 | 300 | 300 |
| Electron exposure (e<sup>-</sup>/Å<sup>2</sup>) | 70 | 70 | 70 |
| Defocus range (μm) | -0.5 to -2.5 | -0.5 to -2.5 | -0.5 to -2.5 |
| Pixel size (Å) | 0.649 | 0.649 | 0.649 |
| # of Movies | 5,580 | 5,498 | 4,777 |
| Initial particle images (no.) | 1,005,231 | 633,138 | 328,914 |
| Final particle images (no.) | 74,671 | 46,329 | 95,511 |
| Symmetry imposed | C2 | C2 | C2 |
| Map resolution (Å) | 2.9 | 2.9 | 2.8 |
| FSC threshold | 0.143 | 0.143 | 0.143 |
| Map sharpening *B* factor | -101.6 | -91.1 | -115.5 |
| ***Refinement*** |  |  |  |
| Initial model used | 4wwr, 7ru9, and<br>AlphaFold3 model | *HsGet3–Get4/Bag6*<br>+ Retro-2 model | *HsGet3–Get4/Bag6*<br>+ Retro-2 model |
| ***Model composition:*** |  |  |  |
| Non-hydrogen atoms | 9895 | 9895 | 9895 |
| Protein residues | 1232 | 1232 | 1232 |
| Ligands | ATP: 2<br>MG: 2<br>ZN: 1<br>HOH: 2<br>Retro-2 (R2): 2 | ATP: 2<br>MG: 2<br>ZN: 1<br>HOH: 2<br>DHQZ36.1 (361): 2 | ATP: 2<br>MG: 2<br>ZN: 1<br>HOH: 2<br>Retro-1 (R1): 2 |
| ***B factors:*** |  |  |  |
| Protein (Å<sup>2</sup>) | 110.99 | 48.79 | 58.78 |
| Ligand (Å<sup>2</sup>) | 67.95 | 32.03 | 42.59 |
| Water (Å<sup>2</sup>) | 50.03 | 13.64 | 9.10 |
| ***R.m.s. deviations:*** |  |  |  |
| Bond lengths (Å) | 0.003 | 0.002 | 0.003 |
| Bond angles (°) | 0.566 | 0.495 | 0.533 |
| ***Validation*** |  |  |  |
| MolProbity score | 1.33 | 1.44 | 1.27 |
| Clashscore | 6.04 | 8.13 | 5.09 |
| Poor rotamers (%) | 0.00 | 0.00 | 0.00 |
| ***Ramachandran plot:*** |  |  |  |
| Favored (%) | 99.36 | 98.47 | 98.79 |
| Allowed (%) | 0.64 | 1.53 | 1.21 |
| Outliers (%) | 0.00 | 0.00 | 0.00 |

## References and Notes

1. H. R. B. Pelham, L. M. Roberts, J. M. Lord, Toxin entry: how reversible is the secretory pathway? Trends in Cell Biology 2, 183–185 (1992).

2. J. S. Bonifacino, R. Rojas, Retrograde transport from endosomes to the trans-Golgi network. Nat Rev Mol Cell Biol 7, 568–579 (2006).

3. R. A. Spooner, D. C. Smith, A. J. Easton, L. M. Roberts, J. M. Lord, Retrograde transport pathways utilised by viruses and protein toxins. Virol J 3, 26 (2006).

4. R. Bingham, H. McCarthy, N. Buckley, Exploring Retrograde Trafficking: Mechanisms and Consequences in Cancer and Disease. Traffic 25, e12931 (2024).

5. M. Holwerda, P. V’kovski, M. Wider, V. Thiel, R. Dijkman, Identification of an Antiviral Compound from the Pandemic Response Box that Efficiently Inhibits SARS-CoV-2 Infection In Vitro. Microorganisms 8, 1872 (2020).

6. W. Dai, Y. Wu, J. Bi, X. Lu, A. Hou, Y. Zhou, B. Sun, W. Kong, J. Barbier, J.-C. Cintrat, F. Gao, D. Gillet, W. Su, C. Jiang, Antiviral effects of Retro-2cycl and Retro-2.1 against Enterovirus 71 in vitro and in vivo. Antiviral Research 144, 311–321 (2017).

7. C. D. S. Nelson, D. W. Carney, A. Derdowski, A. Lipovsky, G. V. Gee, B. O’Hara, P. Williard, D. DiMaio, J. K. Sello, W. J. Atwood, A Retrograde Trafficking Inhibitor of Ricin and Shiga-Like Toxins Inhibits Infection of Cells by Human and Monkey Polyomaviruses. mBio 4, e00729–13 (2013).

8. J. Canton, P. E. Kima, Targeting Host Syntaxin-5 Preferentially Blocks Leishmania Parasitophorous Vacuole Development in Infected Cells and Limits Experimental Leishmania Infections. The American Journal of Pathology 181, 1348–1355 (2012).

9. L. Aeberhard, S. Banhart, M. Fischer, N. Jehmlich, L. Rose, S. Koch, M. Laue, B. Y. Renard, F. Schmidt, D. Heuer, The Proteome of the Isolated Chlamydia trachomatis Containing Vacuole Reveals a Complex Trafficking Platform Enriched for Retromer Components. PLOS Pathogens 11, e1004883 (2015).

10. B. Stechmann, S.-K. Bai, E. Gobbo, R. Lopez, G. Merer, S. Pinchard, L. Panigai, D. Tenza, G. Raposo, B. Beaumelle, D. Sauvaire, D. Gillet, L. Johannes, J. Barbier, Inhibition of Retrograde Transport Protects Mice from Lethal Ricin Challenge. Cell 141, 231–242 (2010).

11. H. Abdelkafi, A. Michau, V. Pons, F. Ngadjeua, A. Clerget, L. Ait Ouarab, D.-A. Buisson, D. Montoir, L. Caramelle, D. Gillet, J. Barbier, J.-C. Cintrat, Structure–Activity Relationship Studies of Retro-1 Analogues against Shiga Toxin. J. Med. Chem. 63, 8114–8133 (2020).

12. J. G. Park, J. N. Kahn, N. E. Tumer, Y.-P. Pang, Chemical Structure of Retro-2, a Compound That Protects Cells against Ribosome-Inactivating Proteins. Sci Rep 2, 631 (2012).

13. N. Gupta, V. Pons, R. Noël, D.-A. Buisson, A. Michau, L. Johannes, D. Gillet, J. Barbier, J.-C. Cintrat, (S)-N-Methyldihydroquinazolinones are the Active Enantiomers of Retro-2 Derived Compounds against Toxins. ACS Med. Chem. Lett. 5, 94–97 (2014).

14. E. Craig, C.-E. Huyghues-Despointes, C. Yu, E. L. Handy, J. K. Sello, P. E. Kima, Structurally optimized analogs of the retrograde trafficking inhibitor Retro-2cycl limit Leishmania infections. PLoS Negl Trop Dis 11, e0005556–e0005556 (2017).

15. D. W. Carney, C. D. S. Nelson, B. D. Ferris, J. P. Stevens, A. Lipovsky, T. Kazakov, D. DiMaio, W. J. Atwood, J. K. Sello, Structural optimization of a retrograde trafficking inhibitor that protects cells from infections by human polyoma- and papillomaviruses. Bioorganic & Medicinal Chemistry 22, 4836–4847 (2014).

16. M. E. Nonnenmacher, J.-C. Cintrat, D. Gillet, T. Weber, Syntaxin 5-Dependent Retrograde Transport to the trans-Golgi Network Is Required for Adeno-Associated Virus Transduction. Journal of Virology 89, 1673–1687 (2015).

17. G. Sivan, A. S. Weisberg, J. L. Americo, B. Moss, Retrograde Transport from Early Endosomes to the trans-Golgi Network Enables Membrane Wrapping and Egress of Vaccinia Virus Virions. J Virol 90, 8891–8905 (2016).

18. K. Harrison, I. R. Haga, T. Pechenick Jowers, S. Jasim, J.-C. Cintrat, D. Gillet, T. Schmitt-John, P. Digard, P. M. Beard, Vaccinia Virus Uses Retromer-Independent Cellular Retrograde Transport Pathways To Facilitate the Wrapping of Intracellular Mature Virions during Virus Morphogenesis. Journal of Virology 90, 10120–10132 (2016).

19. L. Cruz, N. T. Streck, K. Ferguson, T. Desai, D. H. Desai, S. G. Amin, N. J. Buchkovich, Potent Inhibition of Human Cytomegalovirus by Modulation of Cellular SNARE Syntaxin 5. J Virol 91, e01637–16 (2017).

20. O. Shtanko, Y. Sakurai, A. N. Reyes, R. Noël, J.-C. Cintrat, D. Gillet, J. Barbier, R. A. Davey, Retro-2 and its dihydroquinazolinone derivatives inhibit filovirus infection. Antiviral Research 149, 154–163 (2018).

21. X. Zhao, H. Li, J. Li, K. Liu, B. Wang, Y. Wang, X. Li, W. Zhong, Novel small molecule retrograde transport blocker confers post-exposure protection against ricin intoxication. Acta Pharmaceutica Sinica B 10, 498–511 (2020).

22. L. Priyamvada, P. Alabi, A. Leon, A. Kumar, S. Sambhara, V. A. Olson, J. K. Sello, P. S. Satheshkumar, Discovery of Retro-1 Analogs Exhibiting Enhanced Anti-vaccinia Virus Activity. Front Microbiol 11, 603 (2020).

23. E. Craig, A. Calarco, R. Conte, V. Ambrogi, G. G. d’Ayala, P. Alabi, J. K. Sello, P. Cerruti, P. E. Kima, Thermoresponsive Copolymer Nanovectors Improve the Bioavailability of Retrograde Inhibitors in the Treatment of Leishmania Infections. Front. Cell. Infect. Microbiol. 11, 702676 (2021).

24. A. Le Rouzic, J. Fix, R. Vinck, S. Kappler-Gratias, R. Volmer, F. Gallardo, J.-F. Eléouët, M. Keck, J.-C. Cintrat, J. Barbier, D. Gillet, M. Galloux, A New Derivative of Retro-2 Displays Antiviral Activity against Respiratory Syncytial Virus. International Journal of Molecular Sciences 25, 415 (2024).

25. V. Nicolas, V. Lievin-Le Moal, Small Trafficking Inhibitor Retro-2 Disrupts the Microtubule-Dependent Trafficking of Autophagic Vacuoles. Front. Cell Dev. Biol. 8, 464 (2020).

26. S. Madera, F. Izzo, M. F. Chervo, A. Dupont, V. A. Chiauzzi, S. Bruni, E. Petrillo, S. S. Merin, M. De Martino, D. Montero, C. Levit, G. Lebersztein, F. Anfuso, A. Roldán Deamicis, M. F. Mercogliano, C. J. Proietti, R. Schillaci, P. V. Elizalde, R. I. Cordo Russo, Halting ErbB-2 isoforms retrograde transport to the nucleus as a new theragnostic approach for triple-negative breast cancer. Cell Death Dis 13, 447 (2022).

27. D. W. Morgens, C. Chan, A. J. Kane, N. R. Weir, A. Li, M. M. Dubreuil, C. K. Tsui, G. T. Hess, A. Lavertu, K. Han, N. Polyakov, J. Zhou, E. L. Handy, P. Alabi, A. Dombroski, D. Yao, R. B. Altman, J. K. Sello, V. Denic, M. C. Bassik, Retro-2 protects cells from ricin toxicity by inhibiting ASNA1-mediated ER targeting and insertion of tail-anchored proteins. eLife 8, e48434 (2019).

28. U. Kutay, E. Hartmann, T. A. Rapoport, A class of membrane proteins with a C-terminal anchor. Trends in Cell Biology 3, 72–75 (1993).

29. J. W. Chartron, W. M. Clemons, C. J. Suloway, The complex process of GETting tail-anchored membrane proteins to the ER. Current Opinion in Structural Biology 22, 217–224 (2012).

30. M. Amessou, A. Fradagrada, T. Falguières, J. M. Lord, D. C. Smith, L. M. Roberts, C. Lamaze, L. Johannes, Syntaxin 16 and syntaxin 5 are required for efficient retrograde transport of several exogenous and endogenous cargo proteins. Journal of Cell Science 120, 1457–1468 (2007).

31. S. Norlin, V. S. Parekh, P. Naredi, H. Edlund, Asna1/TRC40 Controls β-Cell Function and Endoplasmic Reticulum Homeostasis by Ensuring Retrograde Transport. Diabetes 65, 110–119 (2015).

32. P. T. Linders, C. van der Horst, M. ter Beest, G. van den Bogaart, Stx5-Mediated ER-Golgi Transport in Mammals and Yeast. Cells 8, 780 (2019).

33. P. T. A. Linders, E. C. F. Gerretsen, A. Ashikov, M.-A. Vals, R. de Boer, N. H. Revelo, R. Arts, M. Baerenfaenger, F. Zijlstra, K. Huijben, K. Raymond, K. Muru, O. Fjodorova, S. Pajusalu, K. Õunap, M. ter Beest, D. Lefeber, G. van den Bogaart, Congenital disorder of glycosylation caused by starting site-specific variant in syntaxin-5. Nat Commun 12, 6227 (2021).

34. A. Forrester, S. J. Rathjen, M. Daniela Garcia-Castillo, C. Bachert, A. Couhert, L. Tepshi, S. Pichard, J. Martinez, M. Munier, R. Sierocki, H.-F. Renard, C. Augusto Valades-Cruz, F. Dingli, D. Loew, C. Lamaze, J.-C. Cintrat, A. D. Linstedt, D. Gillet, J. Barbier, L. Johannes, Functional dissection of the retrograde Shiga toxin trafficking inhibitor Retro-2. Nature Chemical Biology 16, 327–336 (2020).

35. J. Rivera-Monroy, L. Musiol, K. Unthan-Fechner, Á. Farkas, A. Clancy, J. Coy-Vergara, U. Weill, S. Gockel, S.-Y. Lin, D. P. Corey, T. Kohl, P. Ströbel, M. Schuldiner, B. Schwappach, F. Vilardi, Mice lacking WRB reveal differential biogenesis requirements of tail-anchored proteins in vivo. Sci Rep 6, 39464 (2016).

36. J. Casson, M. McKenna, S. Haßdenteufel, N. Aviram, R. Zimmerman, S. High, Multiple pathways facilitate the biogenesis of mammalian tail-anchored proteins. J Cell Sci 130, 3851–3861 (2017).

37. M. Y. Fry, V. Najdrová, A. O. Maggiolo, S. M. Saladi, Pavel Doležal, W. M. Clemons, Structurally derived universal mechanism for the catalytic cycle of the tail-anchored targeting factor Get3. Nat Struct Mol Biol, doi: 10.1038/s41594-022-00798-4 (2022).

38. A. F. A. Keszei, M. C. J. Yip, T.-C. Hsieh, S. Shao, Structural insights into metazoan pretargeting GET complexes. Nature Structural & Molecular Biology 28, 1029–1037 (2021).

39. U. S. Chio, S. Chung, S. Weiss, S. Shan, A protean clamp guides membrane targeting of tail-anchored proteins. Proceedings of the National Academy of Sciences 114, E8585–E8594 (2017).

40. H. Cho, Y. Liu, S. Chung, S. Chandrasekar, S. Weiss, S. Shan, Dynamic stability of Sgt2 enables selective and privileged client handover in a chaperone triad. Nat Commun 15, 134 (2024).

41. B. E. Zalisko, C. Chan, V. Denic, R. S. Rock, R. J. Keenan, Tail-Anchored Protein Insertion by a Single Get1/2 Heterodimer. Cell Reports 20, 2287–2293 (2017).

42. M. A. McDowell, M. Heimes, F. Fiorentino, S. Mehmood, Á. Farkas, J. Coy-Vergara, D. Wu, J. R. Bolla, V. Schmid, R. Heinze, K. Wild, D. Flemming, S. Pfeffer, B. Schwappach, C. V. Robinson, I. Sinning, Structural Basis of Tail-Anchored Membrane Protein Biogenesis by the GET Insertase Complex. Molecular Cell 80, 72–86.e7 (2020).

43. S. Shao, M. C. Rodrigo-Brenni, M. H. Kivlen, R. S. Hegde, Mechanistic basis for a molecular triage reaction. Science 355, 298–302 (2017).

44. M. E. Rome, M. Rao, W. M. Clemons, S. Shan, Precise timing of ATPase activation drives targeting of tail-anchored proteins. Proceedings of the National Academy of Sciences 110, 7666–7671 (2013).

45. A. N. Barlow, M. S. Manu, S. M. Saladi, P. T. Tarr, Y. Yadav, A. M. M. Thinn, Y. Zhu, A. D. Laganowsky, W. M. Clemons, S. Ramasamy, Structures of Get3d reveal a distinct architecture associated with the emergence of photosynthesis. Journal of Biological Chemistry 0 (2023).

46. R. Benarroch, J. M. Austin, F. Ahmed, R. L. Isaacson, The roles of cytosolic quality control proteins, SGTA and the BAG6 complex, in disease. Adv Protein Chem Struct Biol 114, 265–313 (2019).

47. J.-Y. Mock, Y. Xu, Y. Ye, W. M. Clemons, Structural basis for regulation of the nucleo-cytoplasmic distribution of Bag6 by TRC35. Proc. Natl. Acad. Sci. U.S.A. 114, 11679–11684 (2017).

48. H. B. Gristick, M. Rao, J. W. Chartron, M. E. Rome, S. Shan, W. M. Clemons, Crystal structure of ATP-bound Get3–Get4–Get5 complex reveals regulation of Get3 by Get4. Nat Struct Mol Biol 21, 437–442 (2014).

49. S. Mukhopadhyay, K. Sinha, S. Chakrabarty, Harnessing Allostery to Modulate Protein–Protein Interactions: From Function to Therapeutic Innovations. Journal of Molecular Biology 437, 169382 (2025).

50. D. Granados-Villanueva, A. Rossow, K. H. Kim, Get4/5-mediated remodeling of Get3’s substrate-binding chamber: Insights into tail-anchored protein targeting by the GET pathway. J Biol Chem 301, 110667 (2025).

51. S. Vajda, D. Beglov, A. E. Wakefield, M. Egbert, A. Whitty, Cryptic binding sites on proteins: definition, detection, and druggability. Current Opinion in Chemical Biology 44, 1–8 (2018).

52. R. Noel, N. Gupta, V. Pons, A. Goudet, M. D. Garcia-Castillo, A. Michau, J. Martinez, D.-A. Buisson, L. Johannes, D. Gillet, J. Barbier, J.-C. Cintrat, N-Methyldihydroquinazolinone Derivatives of Retro-2 with Enhanced Efficacy against Shiga Toxin. J. Med. Chem. 56, 3404–3413 (2013).

53. M. Konstantinidou, M. R. Arkin, Molecular glues for protein-protein interactions: progressing towards a new dream. Cell Chem Biol 31, 1064–1088 (2024).

54. M. L. Repity, R. C. E. Deutscher, F. Hausch, Nondegradative Synthetic Molecular Glues Enter the Clinic. ChemMedChem 20, e202500048 (2025).

55. J. M. Sasso, R. Tenchov, D. Wang, L. S. Johnson, X. Wang, Q. A. Zhou, Molecular Glues: The Adhesive Connecting Targeted Protein Degradation to the Clinic. Biochemistry 62, 601–623 (2023).

56. R. Nussinov, How do dynamic cellular signals travel long distances? Mol Biosyst 8, 22–26 (2012).

57. G. M. Süel, S. W. Lockless, M. A. Wall, R. Ranganathan, Evolutionarily conserved networks of residues mediate allosteric communication in proteins. Nat Struct Mol Biol 10, 59–69 (2003).

58. Y. Ohashi, H. Iijima, N. Yamaotsu, K. Yamazaki, S. Sato, M. Okamura, K. Sugimoto, S. Dan, S. Hirono, T. Yamori, AMF-26, a Novel Inhibitor of the Golgi System, Targeting ADP-ribosylation Factor 1 (Arf1) with Potential for Cancer Therapy. J Biol Chem 287, 3885–3897 (2012).

59. J. B. Saenz, W. J. Sun, J. W. Chang, J. Li, B. Bursulaya, N. S. Gray, D. B. Haslam, Golgicide A reveals essential roles for GBF1 in Golgi assembly and function. Nat Chem Biol 5, 157–165 (2009).

60. B. Y. Kaudeer, J. M. Kirsh, K. Mitachi, J. M. Ochoa, M.-T. Soroush-Pejrimovsky, Y. E. Li, V. N. Nguyen, M. Kurosu, W. M. Clemons, Structures of bacterial and human phosphoglycosyltransferases bound to a common inhibitor inform selective therapeutics. bioRxiv [Preprint] (2025). 10.64898/2025.12.16.694696.

61. R. Nussinov, C.-J. Tsai, Allostery in disease and in drug discovery. Cell 153, 293–305 (2013).

62. P. Martin-Malpartida, C. Torner, A. Martinez, M. J. Macias, TPPU_DSF: A Web Application to Calculate Thermodynamic Parameters Using DSF Data. J Mol Biol 436, 168519 (2024).

63. A. Punjani, J. L. Rubinstein, D. J. Fleet, M. A. Brubaker, cryoSPARC: algorithms for rapid unsupervised cryo-EM structure determination. Nat Methods 14, 290–296 (2017).

64. T. Bepler, A. Morin, M. Rapp, J. Brasch, L. Shapiro, A. J. Noble, B. Berger, Positive-unlabeled convolutional neural networks for particle picking in cryo-electron micrographs. Nat Methods 16, 1153–1160 (2019).

65. J. Jumper, R. Evans, A. Pritzel, T. Green, M. Figurnov, O. Ronneberger, K. Tunyasuvunakool, R. Bates, A. Žídek, A. Potapenko, A. Bridgland, C. Meyer, S. A. A. Kohl, A. J. Ballard, A. Cowie, B. Romera-Paredes, S. Nikolov, R. Jain, J. Adler, T. Back, S. Petersen, D. Reiman, E. Clancy, M. Zielinski, M. Steinegger, M. Pacholska, T. Berghammer, S. Bodenstein, D. Silver, O. Vinyals, A. W. Senior, K. Kavukcuoglu, P. Kohli, D. Hassabis, Highly accurate protein structure prediction with AlphaFold. Nature 596, 583–589 (2021).

66. T. D. Goddard, C. C. Huang, E. C. Meng, E. F. Pettersen, G. S. Couch, J. H. Morris, T. E. Ferrin, UCSF ChimeraX: Meeting modern challenges in visualization and analysis. Protein Science 27, 14–25 (2018).

67. P. V. Afonine, B. K. Poon, R. J. Read, O. V. Sobolev, T. C. Terwilliger, A. Urzhumtsev, P. D. Adams, Real-space refinement in PHENIX for cryo-EM and crystallography. Acta Cryst D 74, 531–544 (2018).

68. P. Emsley, B. Lohkamp, W. G. Scott, K. Cowtan, Features and development of Coot. Acta Cryst D 66, 486–501 (2010).

69. Smart, O.S., Sharff A., Holstein, J., Womack, T.O., Flensburg, C., Keller, P., Paciorek, W., Vonrhein, C. and Bricogne G. (2021) Grade2 version 1.7.1. Cambridge, United Kingdom: Global Phasing Ltd.

70. C. J. Williams, J. J. Headd, N. W. Moriarty, M. G. Prisant, L. L. Videau, L. N. Deis, V. Verma, D. A. Keedy, B. J. Hintze, V. B. Chen, S. Jain, S. M. Lewis, W. B. Arendall III, J. Snoeyink, P. D. Adams, S. C. Lovell, J. S. Richardson, D. C. Richardson, MolProbity: More and better reference data for improved all-atom structure validation. Protein Science 27, 293–315 (2018).

71. J. Pei, N. V. Grishin, PROMALS3D: multiple protein sequence alignment enhanced with evolutionary and 3-dimensional structural information. Methods Mol Biol 1079, 263–271 (2014).

72. J. D. Thompson, D. G. Higgins, T. J. Gibson, CLUSTAL W: improving the sensitivity of progressive multiple sequence alignment through sequence weighting, position-specific gap penalties and weight matrix choice. Nucleic Acids Res 22, 4673–4680 (1994).

73. H. Ashkenazy, S. Abadi, E. Martz, O. Chay, I. Mayrose, T. Pupko, N. Ben-Tal, ConSurf 2016: an improved methodology to estimate and visualize evolutionary conservation in macromolecules. Nucleic Acids Res 44, W344–W350 (2016).

